# Proximity-dependent labeling identifies dendritic cells that drive the tumor-specific CD4^+^ T cell response

**DOI:** 10.1101/2022.10.25.513771

**Authors:** Aleksey Chudnovskiy, Tiago BR Castro, Sandra Nakandakari-Higa, Ang Cui, Chia-Hao Lin, Moshe Sade-Feldman, Brooke K. Phillips, Juhee Pae, Luka Mesin, Juliana Bortolatto, Lawrence D. Schweitzer, Giulia Pasqual, Li-Fan Lu, Nir Hacohen, Gabriel D. Victora

**Affiliations:** Laboratory of Lymphocyte Dynamics, The Rockefeller University, New York, NY, USA; The Francis Crick Institute, London, UK; Broad Institute of MIT and Harvard, Cambridge, MA, USA; Harvard School of Dental Medicine, Harvard University, Boston, MA, USA; Harvard Medical School, Boston, MA, USA; School of Biological Sciences, University of California, San Diego, La Jolla, CA, USA; Moores Cancer Center, University of California, San Diego, La Jolla, CA, USA; Laboratory of Synthetic Immunology, Oncology and Immunology Section, Department of Surgery Oncology and Gastroenterology, University of Padua, Padua, Italy; Veneto Institute of Oncology IOV-IRCCS, Padua, Italy; Center for Cancer Research, Massachusetts General Hospital, Boston, MA, USA

## Abstract

Dendritic cells (DCs) are uniquely capable of transporting tumoral antigens to tumor-draining lymph nodes (tdLNs), and also interact with effector T cells within the tumor microenvironment (TME) itself, mediating both natural antitumor immunity and the response to checkpoint blockade immunotherapy. Using LIPSTIC (Labeling Immune Partnerships by SorTagging Intercellular Contacts)-based single-cell transcriptomics, we identify individual DCs capable of presenting antigen to CD4^+^ T cells in the tdLN as well as inside the tumor microenvironment (TME). Our findings reveal that DCs with similar hyperactivated transcriptional phenotypes interact with helper T cells both within tumors and in the tdLN, and that checkpoint blockade drugs enhance these interactions. These findings show that a relatively small fraction of DCs is responsible for most of the antigen presentation within the tdLN and TME to both CD4^+^ and CD8^+^ tumor-specific T cells and that classical checkpoint blockade enhances CD40-driven DC activation at both sites.

## Introduction

When properly activated, T cells can exert powerful control over cancer, as evidenced by decades of work that culminated in checkpoint-blockade immunotherapies [1–3]. As with other adaptive immune processes, the quality of the T cell response towards a tumor is heavily dependent on the identity and phenotype of the DCs that prime T cells towards tumor-derived antigens in the tumor draining lymph node (tdLN). tdLN DCs are divided into migratory DCs, capable of transporting antigens acquired in the tumor to the tdLN, and resident DCs, which have access only to antigen that arrives to LN through lymphatics or that is handed over to them by their migratory counterparts [4–7]. Both populations can be further subdivided along an orthogonal axis into conventional cDC1 and cDC2 subsets, based on their phenotype and ontogeny [5, 8]. Generally speaking, cDC1s have higher capacity to prime CD8^+^ T cells, whereas migratory cDC2s are best at priming CD4^+^ T cells [9–11], although these distinctions are not absolute [12].

Both cDC1s and cDC2s have been implicated as drivers of the antitumor immune response [9, 13]. Mice lacking cDC1s show enhanced tumor growth concomitant with reduced tumor-specific effector CD8^+^ T cells [11, 14–17]. CD4^+^ T cell priming by cDC2s is suppressed by regulatory T (Treg) cells [18], and Treg depletion unleashes a cDC2-driven CD4^+^ T cell response to the tumor [13]. Moreover, checkpoint inhibitors do not work efficiently in the absence of DCs, highlighting the role of antigen presentation as a major driver of the response to immunotherapy [11, 15, 19]. More recent studies have identified a novel DC state termed “mature DCs enriched in immunoregulatory molecules” (mRegDCs) [20]. DCs in this state have enhanced capacity to take up antigens but reduced ability to prime antitumor T cell responses. mRegDC share many common genes with the novel “cDC3” subset described in humans [21, 22], and further work is needed to clearly delineate the cDC3 and mRegDC states [23].

DCs are also present within the tumor microenvironment (TME), where they can interact with both effector and regulatory T cells, further shaping antitumor immunity. However, given the heterogeneity of DC phenotypes within the TME, as well as the presence at that site of other populations of non-DC antigen-presenting cells (APCs), determining which DC populations contributes to anti-tumor T cell responses remains a challenge [24–26]. Assessing the contribution of specific APC subsets to T cell activity at this site is particularly challenging, because most if not all T cells arriving at the TME have already been primed in the tdLN, and therefore the relative contribution of local TME and distal tdLN APCs to T cell activity within the tumor is difficult to discern [24–27]. To address these types of questions, we previously developed LIPSTIC (Labeling Immune Partnerships by SorTagging Intercellular Contacts), a proximity-based labeling method based on transfer between interacting cells of a labeled substrate detectable by flow-cytometry that allows us to identify and isolate DCs engaged in T cell priming *in vivo* [28]. By combining LIPSTIC with single-cell RNA sequencing (scRNA-seq), we performed interaction-based transcriptomic profiling of the DCs responsible for priming tumor-specific CD4^+^ T cells in the tdLN and of the myeloid cells that engage CD4^+^ T cells in the TME. Our data show that a minor population of DCs, characterized by a hyperactivated transcriptional program, accounted for most CD4^+^ T cell priming in the tdLN, a phenotype that was shared with the DCs that interacted with effector CD4^+^ T cells in the TME. T–DC interaction in both sites were increased by checkpoint blockade with anti-CTLA-4 antibodies. Together, our data indicate that DCs with similar hyperactivated transcriptional phenotypes interact with helper T cells both within tumors and in the tdLN and that checkpoint blockade drugs enhance these interactions.

## Results

### LIPSTIC identifies DCs capable of priming naïve CD4^+^ T cells in response to tumor antigens

LIPSTIC relies on the *S. aureus* transpeptidase sortase A (SrtA) to transfer an injectable biotinylated peptide substrate (biotin-LPETG) between pairs of cells interacting via the CD40L-CD40 pathway. SrtA, fused to the extracellular domain of CD40L expressed on CD4^+^ T cells, captures substrate injected *in vivo* and transfers it onto five N-terminal glycines engineered into the extracellular domain of CD40 (G_5_-CD40) on interacting DCs (**Fig. 1A**). To apply this system to a solid tumor model, we engineered the B16 melanoma cell line to express the OVA_323-339_ (OT-II) peptide (B16^OT-II^; **Fig. S1A**). We then inoculated G_5_-CD40-expressing mice (*Cd40*^G5/G5^) subcutaneously with 10^6^ B16^OT-II^ cells in the flank region to generate a response in the tumor-draining inguinal lymph node (tdLN). At 9 d.p.i., we adoptively transferred into tumor-bearing mice 3 x 10^5^ SrtA-expressing OT-II T cells (either carrying the conditional *Cd40lg*^SrtAv1^ allele crossed to Cd4-Cre or carrying a new, constitutive *Cd40lg*^SrtAv2^ allele, see methods). We performed LIPSTIC labeling of the tdLN by local injection of biotinylated LPETG substrate at 10-12 h after T cell transfer (**Fig. 1B**), a point at which T cell interaction with DCs is exclusively cognate [28]. This system allows transferred OT-II T cells to serve as “reporters” that specifically identify and label the subset of DCs capable of priming naïve CD4^+^ T cells at that time point.

**Figure 1.**
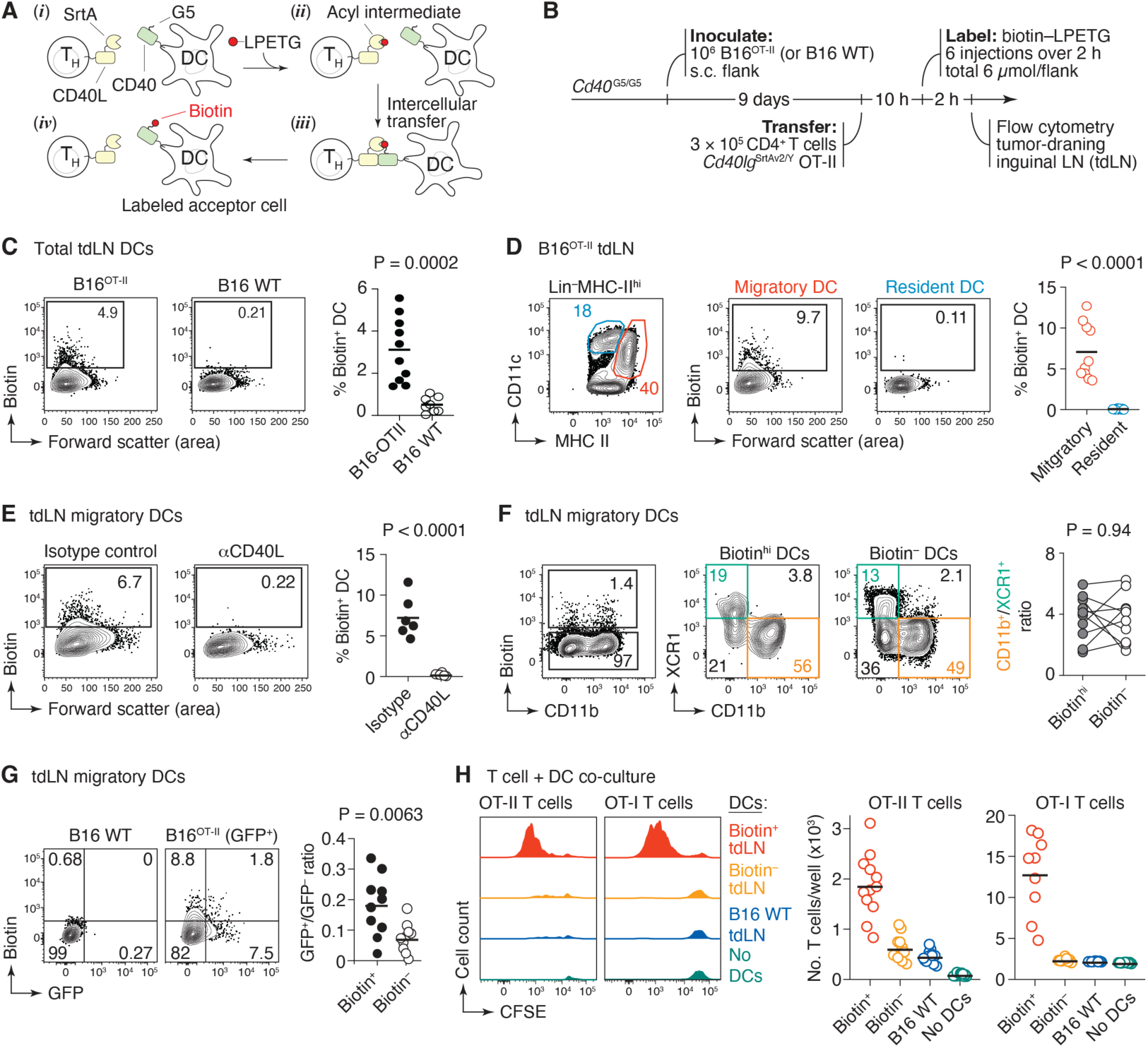
Using LIPSTIC to identify tumor antigen-presenting DCs. **(A)** Schematic of the LIPSTIC system. T cells expressing *S. aureus* transpeptidase sortase A (SrtA) fused to CD40L transfer an injectable biotinylated peptide substrate (biotin-LPETG) onto five N-terminal glycines engineered into the extracellular domain of CD40 (G_5_-CD40) on interacting DCs. **(B)** Experimental setup for panels (C-G). In (C, D, F, G) bilateral tumors were injected. **(C)** Percentage of labeled DCs in mice bearing B16^OTII^ or parental B16 tumors (left) and quantification of data (right) (n = 5, 4 mice per group). **(D)** Gating on resident and migratory DCs in tdLN (left), percentage of labeled migratory and resident DCs in tdLN (center), and quantification of data (n = 5 mice) (right). **(E)** Percentage of labeled DCs in mice treated with isotype control antibody vs. anti-CD40L antibody (left) and quantification of data for (n = 6 mice per group) (right). **(F)** Contour plots (left, center) showing percentage of cDC1 and cDC2 among biotin^+^ and biotin^−^ DCs and quantification of data for (right) ( n = 5 mice). **(G)** Percentage of GFP^+^ and biotin^+^ DCs in tdLN (left) and quantification of GFP^+^ DCs among biotin^+^ and biotin^−^ DCs in tdLN ( n = 5 mice). (C, D, F, G each symbol represents one LN), E each symbol represents one mouse. **(H)** Proliferation of OT-II and OT-I T cells in vitro after 96 h of co-culture with DCs derived from mice carrying B16^OT-II^ or B16^OVA^ tumors, respectively (left) and quantification of T cells per well at the end of the culture period (right). For (H) tdLN were pooled from (n = 6-10 mice) each dot represents one culture well. P-values are for unpaired t test, except (F), where paired t test was used.

This approach revealed that a minor population (approximately 5 to 10%) of DCs acquired the LIPSTIC label in the tdLN of mice bearing B16^OT-II^ tumors but not in controls inoculated with the parental B16 line (B16 WT) (**Fig. 1C**). In agreement with previous findings [10, 28], LIPSTIC labeling in the tdLN was detected exclusively on migratory (CD11c^Int^MHC-II^Hi^) but not resident (CD11c^hi^MHC-II^int^) DCs (**Fig. 1D and S1B**). The specificity of LIPSTIC labeling was confirmed by injection of a blocking antibody to CD40L (clone MR1) 2 h prior to substrate injection, which fully abrogated labeling (**Fig. 1E**). Mixed bone marrow chimera experiments in which a fraction of DCs were deficient for MHC-II confirmed that LIPSTIC captures predominantly interactions driven by antigen presentation in this setting (**Fig. S1C**). Thus, LIPSTIC labeling in the B16 system is antigen-specific and dependent on the CD40L–CD40 interaction. DCs of both XCR1^+^CD11b^−^ (cDC1) and XCR1^−^CD11b^+^ (cDC2) phenotypes (**Fig. 1F**) were labeled by OT-II T cells, at a ratio that corresponded roughly to the total cDC2/cDC1 ratio in each tdLN. Thus, both subsets of DCs are equally capable of priming tumor antigen-specific CD4^+^ T cells, a notion supported by previous findings in the literature [12, 13]. Because our B16^OT-II^ cell line also expresses GFP (**Fig. S1A**), we were able to simultaneously probe DCs for interaction with CD4^+^ T cells and for the extent to which they carry intact tumor-derived proteins. Although there was a statistically significant enrichment in biotin^+^ cells among the GFP^+^ DC population, GFP and LIPSTIC labeling were overall poorly correlated (**Fig. 1G**). Thus, the storage of intact tumor-derived antigen does not overlap substantially with the ability to use this antigen to prime naïve CD4^+^ T cells in vivo, possibly due to the specific kinetics of antigen storage and presentation over time.

To ascertain that biotin^+^ DCs indeed carried the antigen necessary to prime antigen-specific T cells, we set up a miniaturized assay in which 150 tdLN-derived biotin^+^ DCs were co-cultured *ex vivo* with 750 CFSE-labeled OT-II CD4^+^ T cells in the absence of exogenous antigen. This approach showed that biotin^+^ DCs were exclusively capable of driving T cell proliferation above the background levels obtained with DCs derived from mice carrying parental B16 melanomas that lacked the OT-II peptide (**Fig. 1H**). An analogous experiment with OVA-specific CD8^+^ (OT-I) T cells (using a B16 line that we engineered to express a transmembrane version of the full OVA protein, B16^mOVA^) showed that, again, biotin^+^ tdLN DCs were exclusively capable of driving T cell proliferation (**Fig. 1H**). We conclude that LIPSTIC labeling faithfully identifies DCs within the tdLN that are capable of priming both CD4^+^ and CD8^+^ T cells, and that such DCs represent only a minor fraction of all DCs available at that site.

### DCs capable of priming naïve CD4^+^ T cells are in a distinct hyperactivated state

To understand what distinguishes DCs capable of CD4^+^ T cell priming in tdLNs beyond a population-level analysis, we transcriptionally profiled individual biotin^+^ and biotin^−^ migratory DCs obtained from tdLNs at 10 and 15 d.p.i. and control DCs from steady-state iLNs by plate-based single cell mRNA sequencing (scRNA-seq). DCs fell into five transcriptional clusters (**Fig. 2A; Spreadsheet S1**), of which two expressed signatures associated with a cDC1 phenotype (Clusters 1 and 3) and three to a cDC2 phenotype (Clusters 0, 2, and 4) (**Fig. S2A**). Few differences were noted when comparing biotin^−^ migratory DCs sorted from B16^OT-II^ tdLNs or total migratory DCs from steady-state iLNs. On the other hand, biotin^+^ tdLN DCs, especially those with a cDC2 phenotype, were strongly enriched in Cluster 0, which consisted almost exclusively of LIPSTIC-labeled cells (**Fig. 2B and C**). Similar but less pronounced segregation of biotin^+^ and biotin^−^ phenotypes was observed within cDC1 Cluster 3 **(Fig. 2B)**. In both cDC1 and cDC2 populations, LIPSTIC labeling correlated with expression of CD40 target genes [29], as expected given the pathway assayed by the LIPSTIC method (**Fig. S2B**).

**Figure 2.**
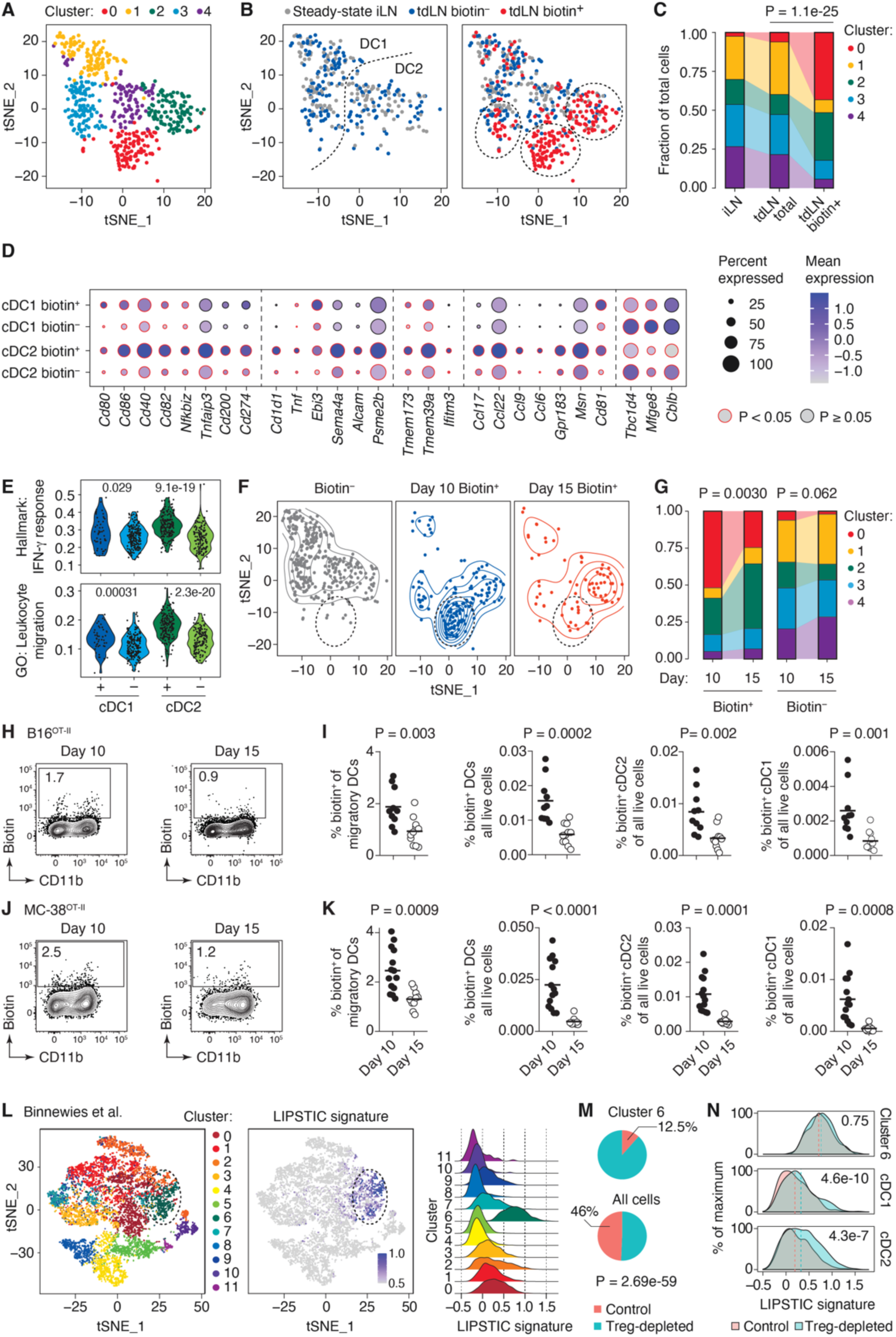
Biotin^+^ DCs represent a transcriptionally distinct DCs state. **(A)** t-distributed stochastic neighbor embedding (t-SNE) scatter plot showing clustering of DCs sorted from tdLN or steady-state iLN. Cells are pooled from 2 steady-state and 2 tumor-bearing mice from 2 independent experiments. **(B)** Distribution of steady-state iLN, biotin^−^ tdLN and biotin^+^ tdLN DCs in the same plot. Dotted lines indicate the approximate location of the cDC1/cDC2 boundary (left) and of biotin^+^ DCs (right). **(C)** The proportion of cells in each transcriptional cluster among steady-state, biotin^−^ and biotin^+^ DCs. **(D)** Expression of genes significantly upregulated in both biotin^+^ cDC1 and cDC2 compared to biotin^−^ cDC1 and cDC2. **(E)** Violin plots show the most upregulated gene signatures in both biotin^+^ cDC1 and cDC2 compared to biotin^−^ cDC1 and cDC2. **(F)** t-SNE contour density plots showing the distribution of biotin^−^ (left) and biotin^+^ DCs at 10 (center) or 15 (right) d.p.i. in tdLN of B16^OT-II^ bearing mice. Dotted circles indicate the approximate location of Cluster 0 (Fig. 2A) **(G)** Distribution of transcriptional clusters from Fig. 2A among biotin^+^ and biotin^−^ DCs from 10 and 15 d.p.i. Data for (F-G) are from the experiment described in Fig. 2A. **(H)** Contour plots show the percentage of biotin^+^ DC in tdLN of mice bearing B16^OT-II^ tumors at 10 (early) or 15 (late) d.p.i. **(I)** Quantification of data as in (H) (n = 10 mice per group from 4 independent experiments, each dot represents one mouse). **(J-K)** as in (H-I) but in mice bearing MC-38^OT-II^ tumors (n = 7 mice per group, with bilateral tumors; each dot represents one tdLN). Experiments in (H-K) used the *Cd40lg*^SrtAv1^ allele. **(L)** t-SNE scatter plot showing clustering (left) and expression of a LIPSTIC+ gene signature (center) among tdLN myeloid cells under control conditions or upon depletion of Treg cells. Right, expression of the LIPSTIC+ signature by cluster. **(M)** Percentage of DCs from control vs. Treg-depleted conditions in Cluster 6 vs. in all cells. **(N)** Expression of the LIPSTIC+ signature in DCs from control or Treg depleted mice in Cluster 6 among total cDC1 and cDC2. Data in (L-N) are from [13]. Pearson’s chi-squared test was used for (C,G and M) Wilcoxon signed-rank test was used for (D, E and N), unpaired t-test was used for (I, K)

Differential gene expression comparing biotin^+^ and biotin^−^ tdLN DCs showed significant modulation of 269 genes (224 upregulated and 45 downregulated). Of these, 136 were commonly modulated in biotin^+^ cDC1 and cDC2 populations and 66 changed exclusively in biotin^+^ cDC2 (**Fig. S2C** and **Spreadsheet S2**). Changes included upregulation of genes encoding for classic markers of DC maturation (*Cd80*, *Cd86*, *Cd40*, *Cd82*) and of nuclear factor (NF)-κB activation (*Nfkbiz*, *Tnfaip3* (A20) [30, 31], typical of an activated DC state, as well as strong upregulation of the inhibitory molecules *Cd200* and *Cd274* (encoding PD-L1) (**Fig. 2D**) [20, 32]. Genes functionally important for T cell priming were also upregulated, including *Cd1d1*, cytokines *Tnf* and *Ebi3*, drivers of DC–T cell interaction such as *Sema4a* and *Alcam* [33–35] and the proteasome activator involved in antigen presentation (*Psme2b*) [36], as were *Ifitm3*, *Tmem39a*, and *Tmem173* (STING), genes important for infectious disease and autoimmunity [37–39]. LIPSTIC-labeled DCs, especially cDC2s, upregulated several genes related to cell migration and microanatomical localization of DCs and T cells, including the chemokines *Ccl17*, *Ccl22*, *Ccl9*, and *Ccl6* [40–44], *Gpr183* (encoding the G-protein-coupled receptor EBI2, a critical guidance receptor that positions DCs at the LN follicle border and splenic bridging channels [45–47], and cell motility regulators such as *Msn* (moesin) and *Cd81* (**Fig. 2D**). On the other hand, biotin^+^ DCs downregulated several genes expressed in anti-inflammatory DCs and/or associated with the induction of T cell tolerance, such as *Tbc1d4*, *Mfge8*, and *Cblb* [48–50] (**Fig. 2D**). Although upregulation of several of these genes was confirmed at the protein level by flow cytometry (**Fig. S2D**), none of the molecules we tested was alone capable of unequivocally distinguishing between biotin^+^ and biotin^−^ DCs. We confirmed findings pertaining to individual genes at the level of entire gene signatures obtained from the “Gene Ontology” and “Hallmark” MSigDB databases [51, 52]. Biotin^+^ cDC1s and especially cDC2s showed higher expression of inflammation/activation signatures as well as of signatures related to cell migration, motility, and regulation of the actin cytoskeleton (**Fig. 2E, Fig. S2E and Data file S3**). Finally, although biotin^+^ DCs expressed higher levels of the “mRegDC” and “cDC3” [20, 53] signatures, overlap between our biotin^+^ LIPSTIC signature and these two gene sets was only moderate, indicating that both programs are related but not identical to that of LIPSTIC-labeled DCs (**Fig. S2F, G**).

Notably, LIPSTIC-labeled DCs in tdLN at the day 15 time point largely lost expression of the Cluster 0 signature (**Fig. 2F and G**), even though they by definition interacted with OT-II T cells via the CD40/CD40L axis. Thus, CD40-mediated interaction with T cells is not sufficient to upregulate the Cluster 0 program in DCs. More importantly, this finding suggested that tumor progression impairs the ability of DCs to prime naïve CD4^+^ T cells in the tdLN. Flow cytometry of LIPSTIC labeling at early (day 10) and late (day 15) time points confirmed this notion, as labeling of migratory cDC1s and cDC2s fell by roughly one-half between 10 and 15 d.p.i. (**Fig. 2H and I**). To extend this observation to a second tumor model, we engineered the MC-38 colon adenocarcinoma cell line [54] to express the OT-II peptide (MC-38^OT-II^; **Fig. S1A**). As with B16 tumors, LIPSTIC labeling of MC-38^OT-II^ by OT-II T cells also decreased with time, again with comparable reductions in cDC1 and cDC2 populations (**Fig. 2J and K**). Thus, DCs capable of priming CD4^+^ T cells decrease in numbers and lose their hyperactivated phenotype as the tumor progresses.

To determine whether DCs with a similar hyperactivated phenotype could be detected in tdLNs in the absence of exogenous T cell transfer, we generated a “LIPSTIC^+^” signature consisting of the 224 genes most highly upregulated in biotin^+^ compared to biotin^−^ DCs (**Spreadsheet S4**). We then applied this signature to a previously published set of single-cell transcriptomes of tdLN myeloid cells obtained at 14 days post-inoculation with B16 melanoma, either in control conditions or after depletion of Treg cells using a *Foxp3*^DTR^ mouse allele. The LIPSTIC signature was expressed predominantly by one group of cells in this dataset (Cluster 6), corresponding to cDC2-phenotype DCs (**Fig. 2L**). Notably, although DCs from control tdLN were also present, the large majority of cells in this group (87.5%) originated from Treg-depleted mice (**Fig. 2M**). Conversely, comparison of DCs from control and Treg-depleted settings showed upregulation of the LIPSTIC signature in cDC2s and to a lesser extent cDC1s, while DCs in Cluster 6 expressed this signature equally regardless of whether they originated in control or Treg-depleted mice (**Fig. 2N**). We conclude DCs with a LIPSTIC-like phenotype can be detected in low numbers in the tdLNs of tumor-bearing mice in the absence of T cell transfer and increase in abundance upon depletion of Treg cells.

Together, our data indicate that DCs actively engaged in the priming of CD4^+^ T cells are in a unique transcriptional state comprising classic features of DC activation, potentially downstream of T cell help itself, as well as increased expression of locomotion and migration genes, suggestive of enhanced ability to colocalize with T cells in LN regions conducive to T cell priming.

### IL-27 produced by priming-competent DCs promotes an effective antitumoral response

Among the most highly upregulated genes in Cluster 0 at 9 d.p.i. were *Ebi3* and *Il27*, which encode for EBI3 and p28—the two subunits of the cytokine IL-27 (**Figs. 3A and S2C**). IL-27 is a pleiotropic cytokine that can take on pro- or anti-inflammatory roles depending on the setting [55, 56], and has been shown to either promote or suppress tumor growth in different experimental models [57, 58]. To investigate the effects of IL-27 in our setting, we treated mice 9 days after subcutaneous inoculation of B16^OT-II^ cells with either a blocking antibody to p28 or an isotype control. One day later, we adoptively transferred OT-II T cells into tumor bearing mice and followed the fate of these cells as they were primed within the tdLN and infiltrated the tumor (**Fig. 3B**). In the tdLN, blocking p28 reduced the ability of OT-II T cells to produce CXCR3 and IFNγ, critical mediators of Th1 effector function [59](**Fig.3C**). In agreement with the role of CXCR3 in promoting effector T cell trafficking to the tumor site [17, 60], p28 blockade reduced OT-II T cell recruitment to the tumor, coinciding with a decrease in production of IFNγ by this population (**Fig. 3D**). Parallel experiments using B16^OVA^ showed that anti-p28 treatment also reduced IFNγ production by OT-I CD8^+^ T cells in both tdLN and tumor, although no difference in expression of CXCR3 or tumor infiltration was observed (**Fig. S3B,C**). To assess the significance to the anti-tumoral response of the loss of T cell effector function upon p28 blockade, we used our engineered B16^mOVA^, which showed delayed growth kinetics in comparison with B16^OT-II^ and was often completely rejected by the host (**Fig. S1A**). Treatment of mice inoculated with B16^mOVA^ with p28-blocking antibody starting at 2 d.p.i. (**Fig. 3E**) resulted in increased tumor size and weight when compared to control-treated mice (**Fig. 3F**). To determine whether IL-27 produced by DCs was required for this antitumor immunity, we inoculated B16^mOVA^ cells into CD11c (Itgax)-cre *Il27*^flox/flox^ mice, in which DCs are deficient in production of IL-27. Although baseline rejection of B16^mOVA^ tumors was less pronounced in control *Il27*^flox/flox^ mice lacking cre recombinase (possibly due to differences in the mouse housing environment at UCSD, where these experiments were performed), tumor growth was still significantly increased in mice lacking DC production of IL-27 (**Fig. 3G**) compared to controls. Thus, IL-27 produced by activated DCs is required for full priming of CD4^+^ T cells and for antitumor immunity in this setting.

**Figure 3.**
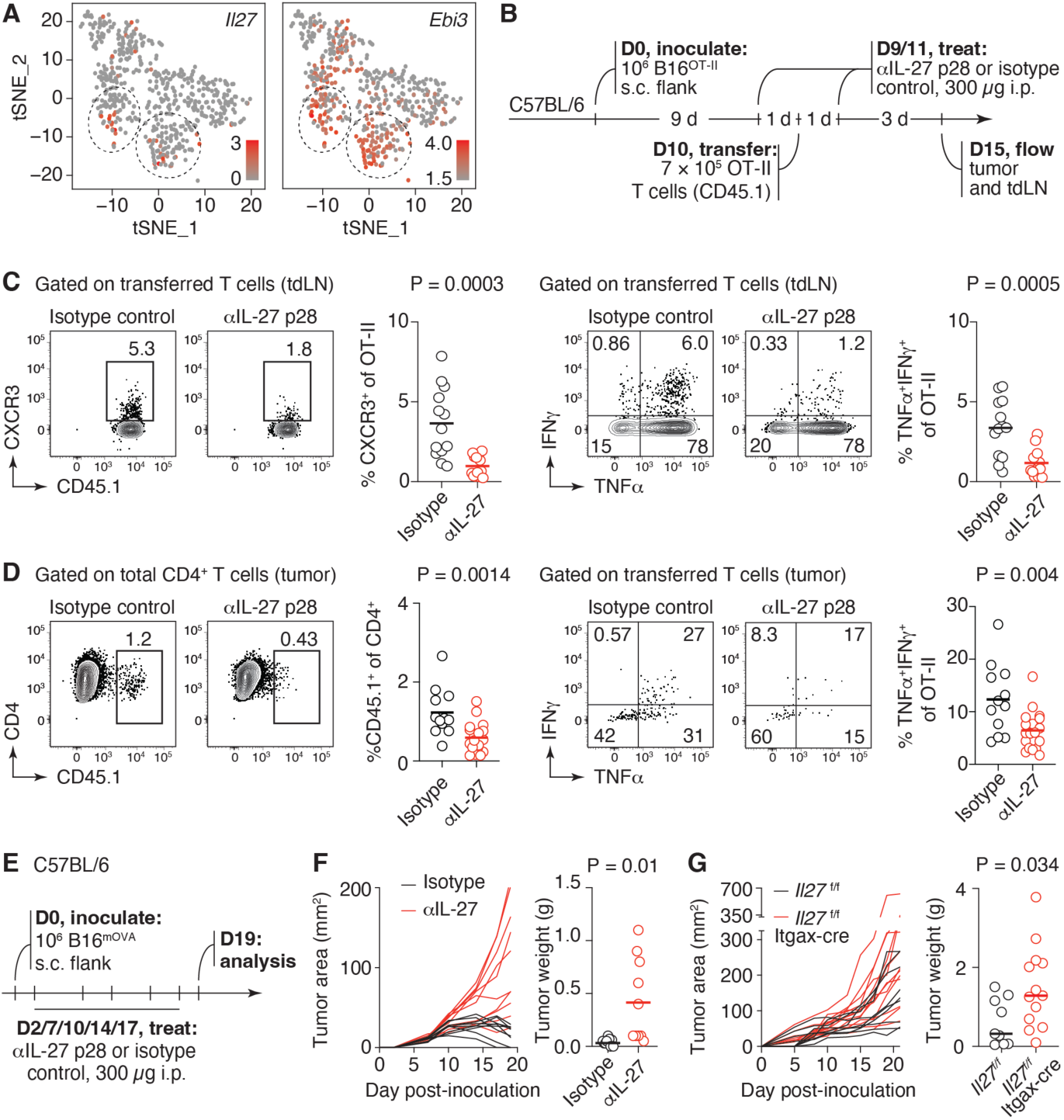
*Il27* promotes an antitumor CD4^+^ T cell response and drives antitumor immunity. **(A)** Expression of *Il27* and *Ebi3* in DCs. t-SNE plot as in Fig. 2A. **(B)** Schematic for IL-27 (p28) blocking in mice bearing B16^OT-II^ tumors. **(C)** Expression of CXCR3^+^ (left) or TNFα and IFNγ (right) among OT-II T cells in tdLN of isotype or anti-p28-treated mice. Graphs summarize data from 2 independent experiments with (n = 13 mice per group.) **(D)** Percentage of OT-II cells among all CD4^+^ T cells (left) and expression of TNFα and IFNγ among OT-II T cells (right) in the tumors of isotype or anti-p28-treated mice. Graphs summarize data from 2 independent experiments with (n = 12 and 18 mice per group). **(E)** Schematic for IL-27 (p28) blocking in mice bearing B16^mOVA^ tumors. (**F**) Tumor growth curves and tumor weights for isotype control and anti-p28-treated groups (n = 9, 10 mice per group). **(G)** Tumor growth curves and tumor weights for *Il27^f/f^* (control) and *Il27^f/f^* Itgax-Cre groups. Data are from 4 independent experiments with n = 9 (control) and n = 13 (*Il27^f/f^* Itgax-Cre). P-values are for unpaired t test.

### Antigen-specific interactions between CD4^+^ T cells and APCs within the TME

Recent studies have shown that DCs can influence antitumor immunity locally within the TME through direct interactions with T cells [25]. Additionally, co-localization of APCs and T cells within the TME in cancer patients correlates positively with responsiveness to immunotherapy [61]. However, the extent to which different APC subsets engage and respond to CD4^+^ T cells under steady-state and immunotherapy conditions is still unclear [24–26]. We measured CD40-dependent interactions between T cells and myeloid cells within the TME by implanting B16^OTII^ tumors into *Cd40*^G5/+^.*Cd40lg*^SrtAv2^ mice, in which all CD40L-expressing endogenous T cells label the CD40-expressing APCs with which they interact (**Fig.4A**). At 10 days post-inoculation, the TME myeloid compartment comprised classical CD11c^+^MHC-II^+^ cDCs and Ly6C^+^MHC-II^hi^ monocytes/monocyte-derived DCs, as well as a large population of F4/80^+^CD11b^+^ macrophages **(Fig.S4A)**. All three populations expressed MHC-II and CD40, and thus had the potential to be labeled by LIPSTIC **(Fig.S4B)**; however, labeling was evident only in a minority of classical DCs and Ly6C^+^MHC-II^hi^ APCs, while labeling of MHC-II^hi^CD40^+^ macrophages was close to background levels (**Fig. 4B**). As in tdLN, cDC1 and cDC2 subsets were labeled in a ratio mirroring the overall cDC2/cDC1 ratio within the TME (**Fig. S4C**). Biotin^+^ TME DCs upregulated the same surface markers as their tdLN counterparts, indicating that T cell-engaged DCs are in an hyperactivated state also within the tumor (**Fig. 4C**). Notably, CD200 upregulation within the TME was found almost exclusively among biotin^+^ cDCs and may thus be useful as a marker for T cell-engaged DCs in WT mice (**Fig. 4C**). Treatment with an anti-MHC-II blocking antibody completely abrogated labeling, suggesting that T cell-APC interactions in the TME are antigen-specific (**Fig. 4D**). This was corroborated by co-culture experiments, which showed that biotin^+^ DC from mice carrying B16^OT-II^ or B16^mOVA^ tumors were more potent drivers of naïve OT-II and OT-I T cell proliferation, respectively, than biotin^−^ DCs from the same tumors (**Fig. 4E**), although TME-resident biotin^−^ DCs were able to drive more substantial proliferation than their counterparts in the tdLN (**Fig. 1H**). Although most APCs were loaded with tumor-derived antigens (asestimated by their GFP fluorescence), there was little if any correlation between GFP fluorescence and LIPSTIC labeling (**Fig. S4D**), indicating that these two measures cannot substitute for each other. We conclude that tumor-infiltrating CD4^+^ T cells interact with TME DCs in an antigen-specific manner, and that this interaction is associated with enhanced DC activation.

**Figure 4.**
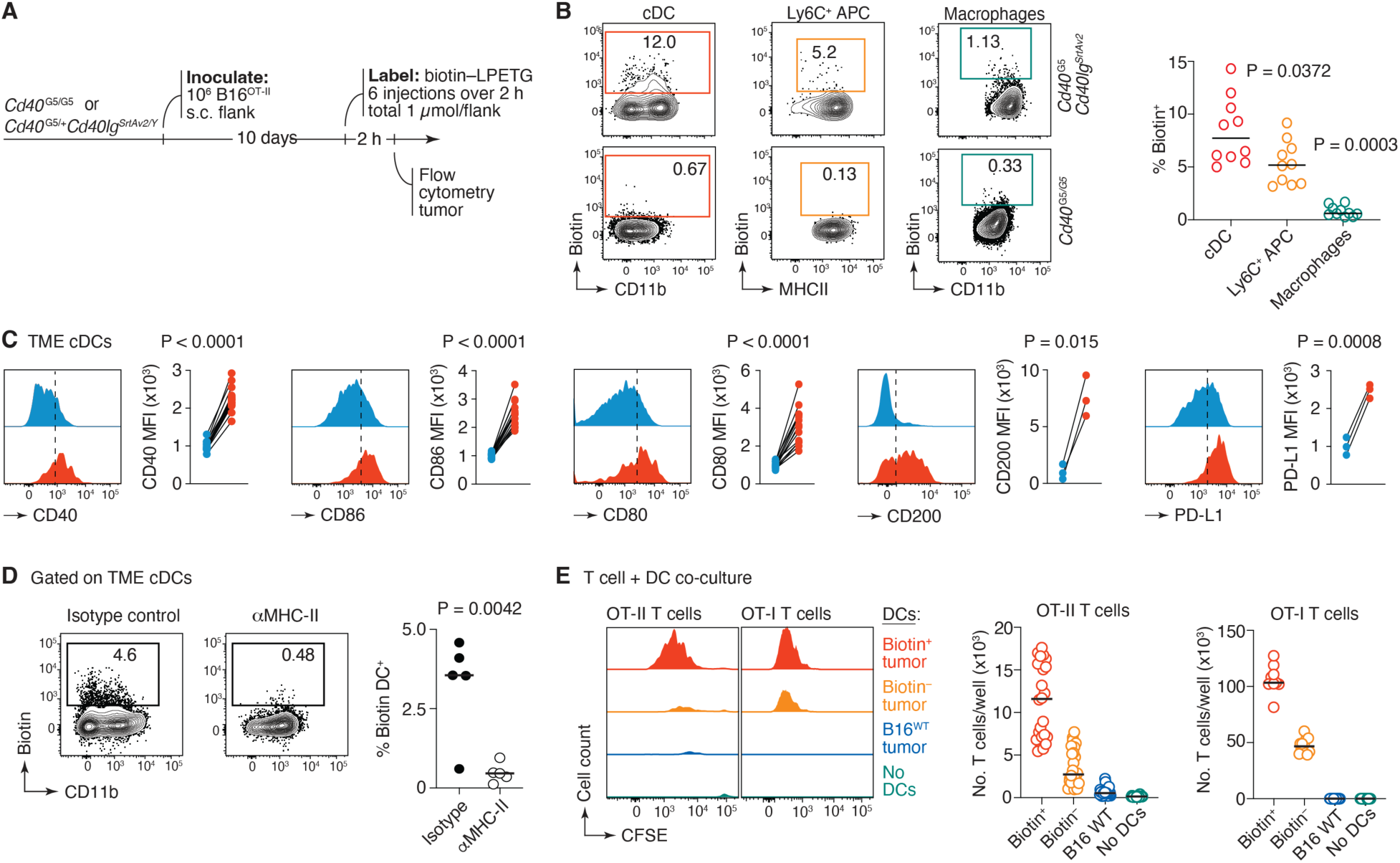
Using LIPSTIC to identify antigen-presenting APCs inside the tumor. **(A)** Experimental setup for panels (B-D). **(B)** Percentage of labeled APCs in (*Cd40*^G5/+^*Cd40lg*^SrtAv2^) mice bearing B16^OTII^ tumors (left) and quantification of data (right) (n = 10 mice, 4 independent experiments) *Cd40*^G5/G5^ mice were used as a baseline for biotin. **(C)** Mean fluorescent intensity (MFI) normalized to the MFI of biotin-negative DC for the indicated molecules. Biotin+(red) and biotin-(blue) DCs within TME (for CD40, CD86, CD80 n = 14 from 4 independent experiments, CD200, PD-L1 n = 3). **(D)** Experimental setup as in (A) except anti-MHC-II or isotype control were injected 2h prior to substrate injection intratumorally (200ug). (Left) Contour plots show percentage of labeled DCs, (right) quantification of labeled DCs (n = 5, from 2 independent experiments). **(E)** Proliferation of OT-II and OT-I T cells in vitro after 96 h of coculture with DCs derived from mice carrying B16^OT-II^ or B16^mOVA^ tumors, respectively (left) and quantification of T cells per well at the end of the culture period (right). For (E), tumors were pooled from (n = 5-9 mice, two independent experiments). Each dot represents one culture well. P-values are for one-way ANOVA(B), paired t-test (C), and unpaired t-test (D).

To better understand the nature of the interactions between myeloid cells and CD4^+^ T cells in the TME, we used droplet-based scRNA-seq combined with an anti-biotin hashtag oligo (HTO)-barcoded antibody [62]. We first performed LIPSTIC labeling in B16^OT-II^ bearing mice as in **Fig. 4A**, except that animals were treated with an isotype control antibody for later comparison with anti-CTLA-4-treated mice (described below). We then sorted TME myeloid cells both as total cells (unenriched), or as a biotin^+^ fraction (enriched) (**Fig. S5A**) at 15 d.p.i. and performed scRNA-seq using the 10x Genomics Chromium platform. Myeloid cells clustered into monocytes/macrophages (Mo/MΦ), cDC1s, cDC2s, and a series of clusters of activated DCs expressing the mRegDC/DC3 signature [20, 53], which we combined into “mRegDC1” and “mRegDC2” clusters based on their proximity to cDC1s and cDC2s and their expression of subtype-specific gene signatures (**Figs. 5A** and **S5B-D** and **Spreadsheet S5**). Whereas LIPSTIC labeling was noted in a small subset of Mo/MΦ, it was most pronounced within the two mRegDC clusters (**Fig. 5B**). Separating cells based on whether they came from the LIPSTIC-enriched or unenriched samples showed that the large size of the Mo/MΦ and mRegDC clusters in the total pool was in part due to the higher abundance of cells with this phenotype among the biotin^+^ population (**Fig. S5E**). Biotin^+^ cells upregulated genes important for antigen presentation (*Psme2*, *Serpinb9*, *H2-DMb2, H2-DMa)* [36, 63, 64] and DC maturation (*Traf1, Stat1)* [65, 66], co-stimulatory molecules (*Cd40*, *Cd1d1, Cd70, Cd200, Cd48)* [67] and chemokines and cytokine important for T cell mediated anti-tumor immunity (*Cxcl9*, *Cxcl16*, *Ebi3*) [17, 58, 68, 69] (**Spreadsheet S5**). Thus, interaction with T cells is associated with an enhanced activation state among myeloid cells in the TME, as it was in the tdLN. Accordingly, all major DC clusters upregulated the genes included in the tdLN_LIPSTIC^+^ signature (**Fig. 5C**), which overlapped substantially, if less than fully, with the corresponding “TME_LIPSTIC^+^” signature obtained by comparing biotin^+^ and biotin^−^ TME DCs (**Fig. S5F**). Thus, as in the tdLN, myeloid cell interaction with T cells in the TME is associated with an enhanced activation state. Plotting trajectories from cDC1 and cDC2 to their adjacent mRegDC subclusters showed that the onset of LIPSTIC labeling either slightly preceded (cDC1) or closely coincided with (cDC2) upregulation of the mRegDC signature in pseudotime, suggesting that CD40L-mediated T cell help may play a role in establishing the mRegDC phenotype (**Fig. 5E,F**). Nevertheless, plotting the expression of the mRegDC signature [20] against that of CD40 target genes [29] revealed two clear populations: an mRegDC x CD40 “diagonal,” comprising mostly cDC1 and cDC2, where expression of both signatures correlated positively, and an mRegDC^hi^CD40^hi^ population comprising cells classified as *bona fide* mRegDC in our clustering (**Fig. 5D**). Thus, although mRegDCs show signs of CD40-mediated activation, CD40 target genes comprised only part of the full mRegDC program, suggesting that other factors, such as antigenic uptake or exposure to cytokines [20], may also be involved in the acquisition of a full mRegDC state.

**Figure 5.**
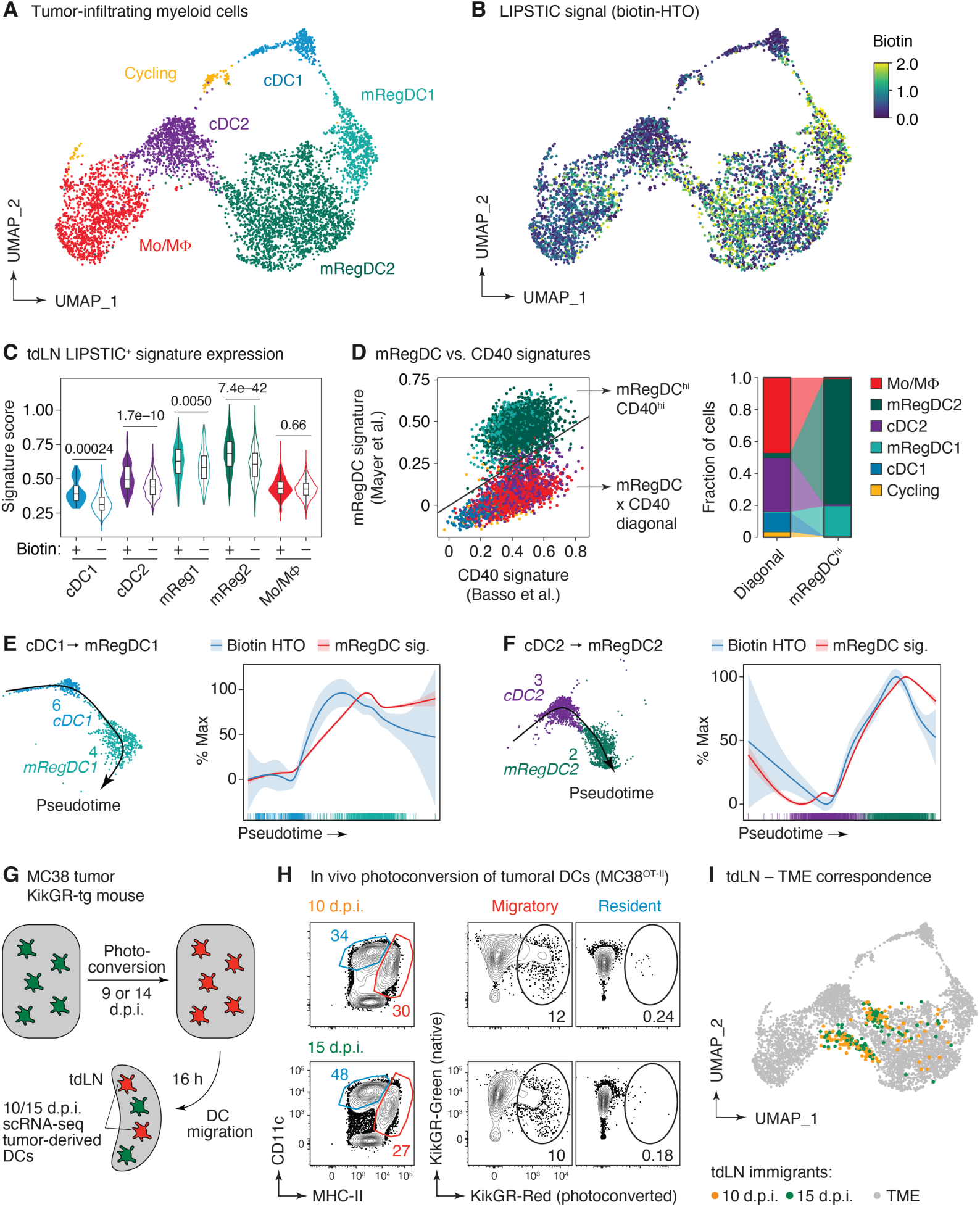
Single-cell interaction-based transcriptomics of APCs within TME. **(A)** Uniform Manifold Approximation and Projection (UMAP) plot showing manually annotated clustering of MHC-II^+^ myeloid cells sorted from the TME at 15 days post-tumor inoculation. See **Fig. S5B** for unsupervised clustering. **(B)** Distribution of LIPSTIC signal (anti-biotin DNA hashtag labeling) in the same cells, in log-normalized counts. **(C)** Expression of the tdLN_LIPSTIC^+^ signature within biotin^+^ and biotin^−^ cells in the indicated populations. P-value is for Wilcoxon signed-rank test. **(D)** Scatter plot showing the expression of CD40 mRegDC gene signatures for the indicated populations of cells (left), quantification of indicated populations in the “mRegDC x CD40 diagonal” and “mRegDC^hi^ CD40^hi^” clusters**. (E)** Pseudotime trajectory showing the transition from cDC1 to mRegDC1 (left) and biotin acquisition and expression of the mRegDC program along that trajectory (right). **(F)** as in (E) but for cDC2. **(G)** Cartoon depicting the tumor photoconversion experiment. **(H)** Percentage of photoconverted migratory and resident DCs in the tdLN at 10 (early, top) and 15 (late, bottom) d.p.i with (n = 6 mice per group; representative plots are shown). **(I)** UMAP showing correspondence between tumor APCs and photoconverted DCs arriving at the tdLN, using the MapQuery function of Seurat.

To gain insight into which of these TME DC populations were most likely to migrate to the tdLN, we used an in-situ photoconversion protocol similar to one previously described in the literature [70] to directly measure the state of DCs arriving from the tumor to the tdLN. We injected mice ubiquitously expressing the photoconvertible *Kikume* green-to-red (KikGR) protein [71, 72] with MC-38^OT-II^ cells, which allow efficient photoactivation due to the lack of the dark pigmentation characteristic of B16 tumors (**Fig. 5G**). Photoactivation of tumors under a 415 nm LED source led to the accumulation of photoconverted migratory, but not resident, DCs in the tdLN as early as 16 h post-photoconversion (**Fig. 5H**), allowing us to isolate DCs immediately after they arrived at the tdLN from the tumor. We performed scRNA-seq profiling of photoconverted DCs sorted from tdLN at 10 and 15 d.p.i. (i.e., 16 h after photoactivation at 9 and 14 d.p.i. respectively; **Fig. 5G**) and used the MapQuery function in the Seurat package (v. 5.0.1) to map photoconverted tdLN DCs to their closest neighbors in the TME 10x dataset (**Fig. 5I**). We found that DCs arriving at the tdLN were most closely related to cells transitioning between the cDC2 and mRegDC2 phenotypes, with fewer cells scattered within the more differentiated mRegDC2 clusters. Plotting the expression of mRegDC and CD40 signatures for photoactivated TME emigrants (**Fig. S5G)** as well as for all tdLN DCs analyzed in the LIPSTIC experiment (**Fig. 2A-G)** showed that they lay mostly on the mRegDC x CD40 diagonal, with no evidence of an additional mRegDC^hi^CD40^hi^ population as found for TME-resident cells (compare **Figs. 5D** and **S5G**). These findings are consistent with a model in which cDC2s exit the TME to migrate to the tdLN prior to fully acquiring the mRegDC signature [73].

### Classical checkpoint inhibitors increase CD40-CD40L interaction axis within the TME

Checkpoint inhibitor-blocking antibodies, such as those targeting CTLA-4 and PD-1, strongly promote antitumoral T cell responses and, as such, have become the key component of tumor immunotherapy. Mechanistically, they are thought to work not only by enhancing the effectiveness of T cell priming in the tumor-draining lymph nodes (tdLN) but also by reviving exhausted T cells within the TME [74, 75]. We first sought to determine whether we could rescue late DC dysfunction in the tdLN (**Fig. 2F-K**) using checkpoint blockade (**Fig. 6A**). Treatment of mice with anti-CTLA-4 led to a marked increase in the number of labeled DCs within the tdLN (**Fig. 6B**), while skewing their composition towards the CD11b^+^ cDC2 subset (**Fig. 6C**). To search for similar effects within the TME, we treated tumor-bearing *Cd40*^G5/+^.*Cd40lg*^SrtAv2^ mice with three injections of anti-CTLA-4 or anti-PD-1 blocking antibodies (or their respective isotype controls) at days 5, 7, and 9 after tumor implantation and performed LIPSTIC labeling and analysis the following day (**Fig. 6D**). Treatment increased the number of biotin^+^ APCs (**Fig. 6E, F**), an effect that was evident among both classical DCs and Ly6C^+^MHC-II^hi^ APCs. PD-1 blockade led to a less pronounced increase among DCs and possibly also Ly6C^+^MHC-II^hi^ APCs (**Fig. S6A**).

**Figure 6.**
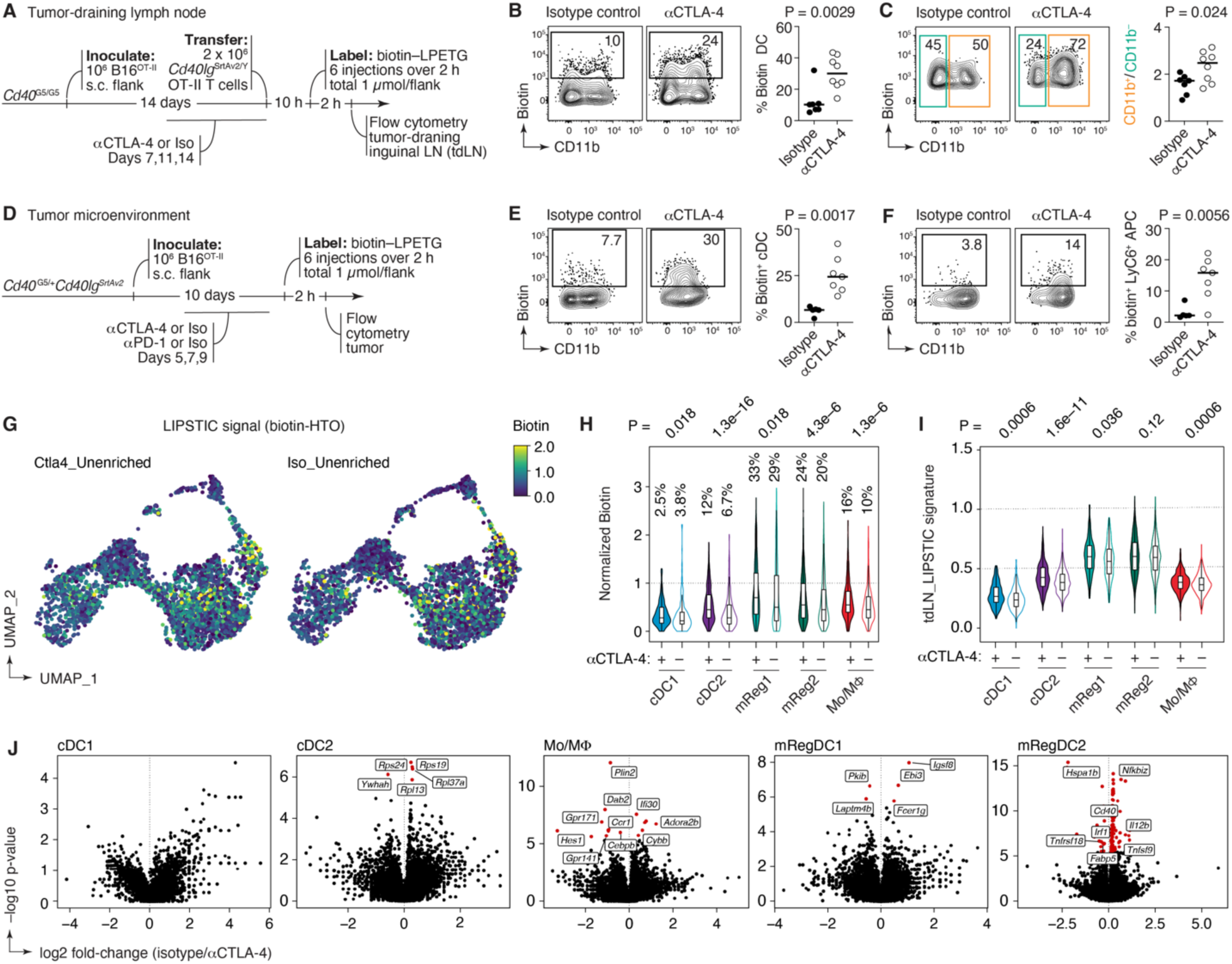
Checkpoint blockade amplifies CD40-CD40L interaction axis in the tdLN and TME. **(A)** Experimental setup for B and C. **(B)** Contour plot and quantification of biotin^+^ DCs in the tdLN in the indicated groups. (n = 7 for iso and n = 8 for CTLA-4, 2 independent experiments) **(C)** Contour plot and quantification of CD11b^+^ and CD11b^−^ DCs among biotin^+^ cells. **(D)** Experimental setup for E and F. **(E)** Contour plot and quantification of biotin^+^ DCs in the TME in the indicated groups. (n = 5 for isotype control and n = 7 for anti-CTLA-4, 2 independent experiments) **(F)** Contour plot and quantification of biotin^+^ Ly6C^+^ APCs in the TME in the indicated groups. P-values were calculated using an unpaired t-test. (**G**) UMAP plot showing the distribution of LIPSTIC signal (anti-biotin DNA hashtag labeling, in log-normalized counts) in anti-CTLA-4 vs isotype control-treated groups (n = 4 for control and n = 3 for anti-CTLA-4). (**H**) Violin plots showing anti-biotin DNA hashtag labeling (LIPSTIC signal) in the indicated populations in anti-CTLA-4 and isotype control groups. Perrcentage of biotin+ cells are given for each violiln. **(I**) Violin plots showing expression of the tdLN_LIPSTIC^+^ signature within anti-CTLA-4 and isotype control-treated groups within the indicated populations. For (H-I) P values were calculated using Wilcoxon signed-rank test. **(J)** Volcano plots showing differentially expressed genes between biotin^+^ APCs from isotype control and anti-CTLA-4-treated mice. Statistically significant genes are colored in red.

Analysis of TME populations by scRNA-seq showed that anti-CTLA-4 treatment increased biotin labeling not only among cDC2s (which showed the largest fold-change in biotin levels) and mRegDCs (which, given their large numbers, accounted for most of the increase in biotin^+^ DCs upon treatment), but also in the Mo/MΦ cluster (**Fig. 6G,H**). Most subsets significantly upregulated the expression of the tdLN_LIPSTIC^+^ signature upon treatment, an increase that was notably more pronounced in cDC populations than in their mRegDC counterparts (**Fig. 6I**). cDC2s additionaly upregulated CD40 target genes (**Fig. S6B**). This pattern suggests that anti-CTLA-4 treatment broadens the acquisition of a hyperactivated state to a larger DC population, but does not result in the establishment of a distinct, checkpoint blockade-specific DC state. Consistent with this, an analysis of differential gene expression comparing only the biotin^+^ TME DCs from anti-CTLA-4 and isotype control-treated mice revealed few significant differences (**Fig. 6J**). Thus, checkpoint blockade, particularly with anti-CTLA-4 antibodies, appears to amplify the extent to which T cells interact with APCs, especially cDC2s, in the TME. This interaction, in turn, leads to increased expression of both tdLN and TME LIPSTIC signatures, especially in less differentiated DC subsets.

## Discussion

A challenge in studying the priming capabilities of DCs has been the heterogeneity in their phenotypes, even within canonical DC populations [48, 76]. Whereas the literature is plentiful in associations between the broader populations of DCs (such as cDC1 and cDC2, or migratory and resident DCs) and the priming of specific types of T cell responses [12, 77], studies that characterize the individual DCs directly engaged in T cell priming are rare [78]. LIPSTIC provides an easy and highly quantitative method to identify and profile T cell-engaged DCs at the individual cell level. As such, LIPSTIC provides degree of specificity to DC research analogous to that afforded to B and T cell studies by antigen tetramers [79, 80].

Our LIPSTIC data revealed that the fraction of all tdLN DCs that actively engage in priming CD4^+^ T cell responses to a subcutaneously implanted tumor is relatively small, on the order of 5-15% of the DCs in the LN even at the early time point peak. These DCs were almost entirely of the CD11c^low^MHC-II^hi^ “migratory” phenotype and were not enriched in cDC1 or cDC2 when compared to the entire LN DC population. In co-culture experiments, tdLN biotin^+^ DCs were exclusively able to prime both CD4^+^ and CD8^+^ T cells. This indicates that the DCs that prime CD8^+^ T cells responses *in vivo* are included within the DC population labeled by CD4^+^ T cells, suggesting substantial overlap between these populations. Thus, CD40L LIPSTIC is unlikely to be missing any DC populations capable of priming exclusively CD8^+^ T cells in the tdLN. Our findings contribute to a series of active discussions in the field regarding the numbers and population-level phenotypes of the DCs that prime the antitumoral T cell response [6, 7, 12, 13, 20, 81].

A potential limitation of LIPSTIC is that it identifies only DCs that engage T cells through the CD40L-CD40 pathway. Most if not all conventional CD4^+^ T cells are thought to express CD40L even in their naïve state [82] and to upregulate its expression upon antigen-driven activation [83]. Conversely, most DCs express CD40, although its expression by resident DCs is generally lower than by DCs with migratory phenotype [84]. Nevertheless, because the CD40L-CD40 axis is critical for DC activation and licensing for cross-presentation [85–88], CD40L LIPSTIC is likely to identify most or all DCs that are potent presenters of immunogenic tumor antigens. This notion is supported by our miniaturized DC culture assays, in which LIPSTIC-negative DCs were unable able to induce proliferation of CD4^+^ or CD8^+^ T cells above background levels, indicating that most antigen-presenting capability is concentrated within the LIPTSIC^+^ DC population.

Interaction-based transcriptional profiling of LIPSTIC-labeled cells showed that priming-capable DCs expressed a unique transcriptional profile that separated them from other DCs in the tdLN. Notably, total DCs from tdLN were indistinguishable transcriptionally from DCs obtained from a naïve iLN, implying that this population of DCs, or at least their prominent role in T cell priming, would be difficult to identify without LIPSTIC. The ability to prime T cells was associated with upregulation of multiple genes important for DC activation and antigen presentation. These included classic DC maturation markers such as *Cd80*, *Cd86*, and *Cd40* itself [89], indicative of partial overlap with the canonical “mature DC” phenotype [90]. However, LIPSTIC-labeled DCs also upregulated multiple genes and signatures associated with cell migration, adhesion, and T cell chemotaxis [91]. Thus, the ability of DCs to access specific microanatomical compartments and to recruit T cells to these areas, previously shown to be functionally important for T cell responses [45–47, 92], is in fact one of the defining characteristics of the DCs capable of engaging T cells *in vivo*. Intersecting our transcriptional data with a much larger scRNA-seq data set of tdLN myeloid cells [13], revealed a cluster of DCs with strong expression of the tdLN_LIPSTIC^+^ signature. A small fraction of this cluster represented DCs found in the tdLN under control non-manipulated conditions—indicating that the tdLN_LIPSTIC^+^ transcriptional state is not a product of our specific setup (e.g., of the adoptive transfer of a relatively large number of tumor antigen-specific T cells). More importantly, however, the large majority of cells in this cluster consisted of DCs that appeared upon ablation of Treg cells using a DTR transgene. These findings suggest that one of the effects of Treg cells, possibly potentiated in the tumor microenvironment, is to prevent DCs from acquiring a LIPSTIC^+^-like state. Of note, the cytokine IL-27 was among the most upregulated gene products in biotin+ DCs. IL-27 is an especially pleiotropic cytokine, which has been shown to prime both tolerogenic and immunogenic T cell responses under different conditions [55–58, 69, 93–96]. In our settings, IL-27 produced by DCs was essential for priming IFNγ production by tumor-specific CD4^+^ and CD8^+^ T cells, recruitment of effector T cells to the tumor site, and control of tumor growth. This result agrees with previous reports in the literature and disagrees with others under similar settings [57, 58]. The reasons for these discrepancies remain unclear and merit further investigation.

Myeloid cells interact with T cells within the tumor microenvironment (TME), shaping anti-tumor responses locally [6, 24, 25]. Quantifying the relative contribution of various antigen-presenting cells (APC) to CD4 T cell stimulation within the TME is challenging, due to the vast heterogeneity of myeloid compartment within tumors [6, 24, 25, 97]. Our data show that DCs and, to a lesser extent, Ly6C^+^ APCs were the primary populations interacting with CD4^+^ T cells within the TME. Despite high CD40 and MHC-II expression, macrophages engaged minimally with T cells, indicating that factors beyond the expression levels of these molecules influence APC-T cell interactions inside the tumor. The ability of DCs to interact with T cells correlated with the expression of classical activation markers, such as CD80, CD86, and CD40, as well as with the expression of the tdLN-derived LIPSTIC^+^ signature. Thus, DCs that engage with T cells in the TME are in a hyperactivated state similar to that found among interactors in the tdLN. Our scRNA-seq data showed that DC labeling correlated strongly with (and possibly preceded in time) expression of the mRegDC program in both cDC1 and cDC2 subsets. This suggests the possibility that, in addition to antigenic capture and IFNγ signaling [20], the mRegDC state may be partially induced by CD40L-mediated T cell help. Additional work will be required to strictly determine whether T cell help drives the mRegDC state or, conversely, it is the acquisition of the mRegDC state that poises DCs to engage with CD4^+^ T cells. Interestingly, when we overlaid the scRNA-seq from photoconverted DCs to our broader 10x dataset, we observed that photoconverted DCs, which emigrated from the tumor to the tdLN, did not fall into the fully differentiated mRegDC cluster present within TME, but rather resembled less mature cDC2s. This suggests an existence of additional signals that trigger the exit of DCs from TME to the tdLN prior to their full maturation. Speculatively, help provided by CD4^+^ T cell to DCs early after their arrival to the TME may drive DC migration to the tdLN, while T cells that remain in the TME may acquire the fully differentiated mRegDC phenotype.

Checkpoint inhibitors have revolutionized cancer therapy, yet the mechanisms underlying their action are not fully resolved [74, 98]. Our findings corroborate a previous study indicating that CTLA-4 treatment increases cDC2 migration to the tdLN [13]. Furthermore, we observed that the amount of interaction between CD4^+^ T cells and DCs is markedly enhanced by checkpoint inhibitors in both the tdLN and the tumor itself. Interestingly, the most significant increase in expression of tdLN_LIPSTIC^+^ signature genes upon anti-CTLA-4 treatment was found among cDC2 and cDC1 subsets, suggesting that checkpoint blockade accelerates the acquisition of hyperactivated DCs state in DCs. Additionally, we observed a significant increase in CD40 signature exclusively in cDC2. These results suggest that CTLA-4 blockade may exert its effects in part by promoting the interaction of CD4^+^ T cells with incompletely matured cDC2s in the TME, triggering their activation and migration to the tdLN. This effect can be aided by increased T cell-cDC interactions in the tdLN, as well as by T cell-mRegDC interactions directly within the TME.

In conclusion, LISPTIC allowed us, for the first time, to identify the individual DCs capable of priming CD4^+^ T cells in tdLNs and engaging with these cells within the TME, and to define transcriptional programs associated with these interactions at both sites. The *in vivo* DC activation signature identified in this work has the potential to be exploited practically to improve responses to tumoral antigens by therapeutic targeting of DCs or their products. In addition, we expect that the LIPSTIC interaction-based transcriptomics platform laid out in this study will be useful for immunologists wishing to identify the DCs that prime CD4^+^ T cell responses in various settings.

## Acknowledgments

We thank A. Kamphorst (Mount Sinai School of Medicine) and C. Brown (Memorial Sloan Kettering Cancer Center) for critical reading of our manuscript; Z. Yin (Nankai University) for *Il27*^flox^ mice; C. Brown for the B16^OVA^ cell line; K. Gordon for cell sorting; the Rockefeller University Genomics Center for RNA sequencing and Comparative Biosciences Center for animal housing; and Rockefeller University employees for their continuous assistance. This work was supported by NIH grant DP1AI144248 and the Hirschl/Weil-Caulier Research Award (G.D.V.), Starr Consortium grant I10-044 (N.H. and G.D.V.) and NIH grants AI108651 and AI163813 (L.-F.L.). Work in the Victora Laboratory is supported by the Robertson Foundation and the he Stavros Niarchos Foundation (SNF) Institute for Global Infectious Disease Research. A.C. was supported by a Damon Runyon Postdoctoral Fellowship. G.D.V. is a Pew-Stewart Scholar.

## Author contributions

A.C. designed and performed most experiments, with assistance from G.P., B.K.P., J.P., and L.M.. S.N.-H. and J.B. generated *Cd40lg*^SrtAv2^ mice. T.B.R.C. and A.C. performed all computational analyses. M. S.-F. and L.D.S. generated single-cell RNA-seq libraries under supervision of N.H. C.-H.L. and L-F.L. designed and performed experiments on *Il27*-deficient mice. N. H. and G.D.V. designed and supervised experiments. G.D.V. and A.C. wrote the manuscript with input from all authors.

## Data availability

Gene expression data are deposited in GEO under accession number GSE275471

## MATERIALS AND METHODS

### Mice

*Cd40*^G5^ mice were generated as described [28] and maintained in our laboratory. Two versions of *Cd40lg*^SrtA^ mice were used, the original conditional version [28] (which we refer to as *Cd40lg*^SrtAv1^) and a new constitutive *Cd40lg*^SrtAv2^ version (see below). For intratumoral experiments, we have generated *Cd40*^G5/+^.*Cd40lg*^SrtAv2^ mice, by crossing *Cd40*^G5^ with constitutive *Cd40lg*^SrtAv2^. Unless indicated in the figure legend, all experiments are done using the constitutive version. C57BL6/J, CD45.1 (B6.SJL Ptprca), *H2*^−/−^ [99], CD4-Cre-transgenic [100], and OT-I TCR transgenic [101] mice were purchased from The Jackson Laboratory (strains 000664, 002014, 003584, 022071, and 0003831, respectively). OT-II TCR-transgenic mice (Y chromosome version) [102] mice were bred and maintained in our laboratory. *Il27*^flox^ [69] and Itgax-cre [103] mice were bred and maintained at UCSD. CAG-KikGR-transgenic mice [71] were a gift from A. Hadjantonakis (Memorial Sloan Kettering Cancer Center). CAG-KikGR-transgenic mice were back-crossed to the C57BL6 background for at least 10 generations at Rockefeller University. All mice were housed in specific pathogen-free conditions, in accordance with institutional guidelines and ethical regulations. All protocols were approved by the Rockefeller University or UCSD Institutional Animal Care and Use Committees. Male and female mice aged 5–12-weeks were used in all experiments.

### Generation of Cd40lg^SrtAv2^ mice

*Cd40lg*^SrtAv2^ mice were generated in our laboratory using the EASI-CRISPR method [104]. Cas9-crRNA-tracrRNA complexes targeting the last exon of the *Cd40lg* locus (GAGTTGGCTTCTCATCTTT) were microinjected along with a single-stranded DNA templates encoding a C-terminal SrtA fusion flanked by 200bp homology arms into the pronuclei of fertilized C57BL6 embryos, which were then implanted into pseudopregnant foster dams. Founder mice were backcrossed to WT C57BL6 mice for at least 5 generations to reduce the probability of transmitting CRISPR-induced off-target mutations.

### Generation of transgenic tumor lines

Constructs were cloned into the pMP71 vector [105], which was modified to express a fluorescent reporter (eGFP) followed by sequence encoding amino acids 323-339 of chicken OVA (eGFP-OT-II) or the *thosea asigna* virus self-cleaving 2A peptide (T2A) [106]followed by the full-length OVA protein (eGFP-OVA). Retroviruses were produced in HEK-293 cells using CaCl2 transfection. Gag-Pol and VSV plasmids were used for virus packaging. Virus-containing supernatant was harvested 48 h after transfection, spun down at 252 XG for 5 min, and filtered through a 0.45 µm filter. B16-OT-II or B16-OVA^TM^ melanoma cell lines or MC38-OT-II colon adenocarcinoma were produced by retroviral transduction with eGFP-OT-II or eGFP-OVA^TM^ constructs. Briefly, viral supernatant from HEK-293 cells was added to B16 or MC-38 cells with 5 mg/ml of polybrene (Sigma-Aldrich H9268) and cells were spun down at 800 XG for 90 minutes at 30 degrees. Viral supernatant was replaced with regular medium 12 h after transfection. 60 h after transfection, eGFP^+^ cells were sorted using a FACSAria II, expanded, and frozen in liquid nitrogen.

### Murine tumor models

All tumor cell lines were grown in DMEM supplemented with 1% L-Glutamine and 1% Penicillin/ Streptomycin. On the day of tumor injection, tumor cells were trypsinized with TrypLE express, washed twice, and resuspended in sterile PBS. Mice were anesthetized and shaved on one or both flanks, and one million of tumor cells were injected in 50 µl of PBS subcutaneously (unilaterally or bilaterally) at a site adjacent to the inguinal lymph. Tumor growth was assessed 2-3 times per week by caliper measurements. Tumor area was calculated by multiplying tumor length and tumor width.

### Adoptive cell transfers

To isolate CD4^+^ or CD8^+^ T cells, spleens were forced through 70 µm filters, red blood cells were lysed with ACK buffer (Lonza), and CD4^+^ or CD8^+^ T cells were isolated using CD4^+^ or CD8^+^ T cells isolation kits (Miltenyi Biotec) as described in the manufacture protocols. Purified cells were injected intravenously in 100 µl of PBS.

### LIPSTIC in vivo

Biotin-aminohexanoic acid**-**LPETGS, C-term amide, 95% purity (biotin-LPTEG) was purchased from LifeTein (custom synthesis) and stock solutions prepared in PBS at 20 mM. For LIPSTIC labeling in vivo, biotin–LPETG was injected subcutaneously into the flank (shaved area proximal to tumor and iLN). 50 μl of 20 mM substrate dissolved in PBS (equivalent to 1 μM/injection) were injected a total of six times (6 μM total) 20 min apart, and iLN were collected 20 min after the last injection, as described previously [28]. Mice were briefly anaesthetized with isoflurane before each injection. For labeling intratumoral APC-T cell interactions, everything was done as above except substrate same amount was injected intratumorally.

### Antibody treatment

For IL-27 blocking experiments, animals were injected intraperitoneally with 300 µg of p28 blocking antibody clone (BioXcell MM27.7B1) or isotype control clone (BioXcell C1.18.4). The regimen for each experiment is described in the corresponding figure legend. For CD40L-blockade experiments, mice were injected intravenously with 200 μg of CD40L-blocking antibody (clone MR-1, BioXCell) two hours prior to the first injection of substrate. For anti-CTLA-4 treatment, animals were injected as shown in the figure with 200ug of clone 9H10 (BioXcell) or corresponding isotype control Syrian Hamster IgG (BioXcell). For anti-PD1 treatment, animals were injected as shown in the figure with 200ug of clone RMP1-14 (BioXcell) or corresponding isotype control rat IgG2a clone 2A3 (BioXcell). For MHC-II blocking experiment mice were injected intratumorally with anti MHC-II blocking antibody (I-A/I-E) (clone M5/114, BioXCell) 2h prior to first substrate administration or isotype rat IgG2b (clone LTF-2, BioXcell).

### Surgery and KikGR photoconversion

Tumor-bearing animals were anaesthetized with isoflurane, shaved, and washed with ethanol and 0.005M iodine solution. The tumor was completely exposed by an incision made on the side distal to the iLN. Photoactivation was performed by exposing the tumor to 415 nM light for three minutes using a custom-built Prizmatix LED source. The incision was then closed using FST autoclips.

### Bone marrow chimeras

WT C57BL6/J mice were lethally irradiated with two doses of 450 rad from an X-ray source, administered five hours apart. Animals were reconstituted by intravenous injection of hematopoietic cells obtained from bone-marrows of donor mice. Chimeric mice were used in experiments 7-12 weeks after irradiation.

### Flow cytometry and cell sorting

iLN were collected and cut into small pieces in microfuge tubes. For digestion, iLN were incubated for 30 min at 37 °C in HBSS (Gibco) supplemented with CaCl_2_, MgCl_2_, and collagenase D (Roche). After digestion, tissue was forced 5 times through a 21 Gauge needle (BD) and filtered through a 70 µm strainer into a 15 ml falcon tube. Tumors were excised, cut into small pieces and digested with collagenase D and DNase I for 1 h at 37 °C in HBSS (Gibco). After digestion, tumors were forced 10 times through a 21 gauge needle (BD) and filtered through a 70 µm strainer into a 50 ml falcon tube. Tumors were spun down and resuspended in 3 ml 10% Percoll/RPMI, which was layered on top of 90% Percoll/RPMI. Cells were spun down for 25 min at 615 XG at 20°C with breaks off. The gradient interface containing leukocytes was collected and washed with 15 ml of PBE buffer. Single-cell suspensions were washed with PBE and incubated at room temperature for 5 min with 1 μg/ml anti-CD16/32 (2.4G2, BioXCell). Cells were stained for surface markers on ice in 96-well plates for 15 min in PBE, using the reagents listed in Table S1. Cells were washed with PBE and stained with Zombie fixable viability dyes (Biolegend) at room temperature for 15 min, then fixed with Cytofix (BD Biosciences). For biotin–LPETG SrtA substrate staining, an anti-biotin–PE antibody (Miltenyi Biotec) was exclusively used as described [28]. Samples were acquired on Symphony, Fortessa or LSR-II flow cytometers or sorted on FACSAriaII or FACSAriaIII cell sorters (BD Biosciences). Data were analyzed using FlowJo v.10.6.2 software.

### In vitro lymph node DC-T cell co-culture

Mice were injected with the indicated tumors. On day 10, animals were injected with substrate as described above, and biotin^+^, biotin^−^, or total migDCs from OVA^−^ tumors (150 DCs for OT-II and 500 DCs for OT-I cultures) were sorted into U-bottom 96-well plates. 750 CD4^+^ OT-II and 2500 CD8^+^ OT-I CFSE-labeled splenic T cells were sorted in the corresponding wells. All wells contained RPMI supplemented with 1% Penicillin/Streptomycin, 1% Sodium Pyruvate, 1% non-essential amino acids, 1% HEPES. For OT-II cell cultures, RPMI media was supplemented with 2% T-stim media with ConA (Avantor). All cells were transferred into FACS tubes and the entire sample was recorded to determine cell numbers.

### In vitro tumor DC-T cell co-culture

Mice were injected with the indicated tumors. On day 10, animals were injected with the substrate intratumorally as described above, and biotin^+^, biotin^−^, or total migDCs from OVA^−^ (700 DCs for OT-II and 700 DCs for OT-I cultures) were sorted into U-bottom 96-well plates. 3500 CD4^+^ OT-II and 3500 CD8^+^ OT-I CFSE-labeled splenic T cells were sorted in the corresponding wells. All wells contained RPMI supplemented with 1% Penicillin/Streptomycin, 1% Sodium Pyruvate, 1% non-essential amino acids, 1% HEPES.

### Library preparation for scRNA-seq and bulk RNA sequencing

Libraries were prepared using the Smart-Seq2 method, as previously described [107]. In brief, RNA from single-sorted cells was extracted using RNAClean XP Solid Phase Reversible Immobilization (SPRI) beads (Beckman Coulter). Extracted RNA was first hybridized using using an RT primer (/5BiosG/AAGCAGTGGTATCAACGCAGAGTACTTTTTTTTTTTTTTTTTTTTTTTTTTTTTTVN), then reverse transcribed into cDNA using a TSO primer (59-AAGCAGTGGTATCAACGCAGAGTACATrGrGrG-39) and RT maxima reverse transcriptase (Thermo Fisher Scientific). Amplification of cDNA was performed using an ISPCR primer (59-AAGCAGTGGTATCAACGCAGAGT39) and KAPA HiFi HotStart ReadyMix (Thermo Fisher Scientific). Amplified cDNA was cleaned up three times using RNAClean XP SPRI beads. cDNA was tagmented using the Nextera XT DNA Library Preparation Kit (Illumina). For each sequencing run, up to four plates were barcoded at a time with Nextera XT Index Kit v2 Sets A–D (Illumina). Dual-barcoded libraries were pooled and sequenced using Illumina Nextseq 550 platform.

### Library preparation for single cell-RNA sequencing of tumor APCs

Single-cell RNA-seq libraries of tumor APCs were prepared by staining cells with oligonucleotide-conjugated antibodies to CD45; MHC-I, and anti-biotin for the LIPSTIC signal detection. Post-sorting, cells were gathered in a microfuge tube containing PBS 0.4% BSA, concentrated by centrifugation, and adjusted to a final volume of 35-40 ul. Viability counts were performed, and cells were immediately processed for library construction using the Chromium platform (10x Genomics), following the manufacturer’s guidelines. The Genomics Core at Rockefeller University carried out the sequencing on an Illumina NovaSeq SP flowcell, aiming for at least 30,000 reads per cell with specific read lengths for each segment of the sequencing process.

### Smart-seq2 transcriptomic analysis

Plate-based scRNA-Seq libraries were processed by applying the STAR aligner for transcriptome alignment, using the GRCm38.p6 mouse genome assembly and GENCODE M20 mouse annotations. Quantification matrices were produced using RSEM. For our single-cell studies, quantification matrices were loaded into the R environment and processed using the Seurat package pipelines [108]. Briefly, cells containing more than 10% of their reads mapped to mitochondrial DNA were filtered out. In addition, we also removed single-cell contaminants that expressed known markers of B-cells, T-cells, and macrophages. In total, 552 cells were selected for downstream analysis for our LIPSTIC experiment and 303 cells for our photoactivation experiment. For both experiments, cells were log normalized, and the top 2,000 variable genes were selected for data scaling and PCA construction. During data scaling, the library size and the mitochondrial percent variables were ‘regressed out’ from the post-normalized data. After PCA construction, the JackStraw algorithm available in Seurat was used to select the most significant principal components. Finally, cells were clustered and visualized in t-SNE space. For signature analysis, the AddModuleScore function was used to determine the average expression of a gene list compared to a background control. Gene set enrichment analysis (GSEA) was performed in the R environment using the fgsea package [109] to determine pathway enrichment. Shortly, fgsea was run using signature sets obtained from MSigDB and used as an input with a pre-ranked list based on log2 fold change calculated from different pairwise comparisons. Signatures were considered enriched if the adjusted p-value had a value of at least 0.05. To determine commonly activated genes in LIPSTIC-positive cDC1 and cDC2 cells, we used a LIPSTIC signature as a pathway and the pre-ranked comparison between cDC1 (positive x negative) and cDC2 (positive x negative). We then compared the two LIPSTIC positive leading edges for matching and exclusive genes. For external data analysis, data was downloaded from the GEO database, accession code: GSE125680 [13]. To apply our LIPSTIC signature, we used the AddModuleScore function available with the Seurat package. We used the top 20 genes enriched in biotin-positive DCs compared to biotin-negative DCs, or the top 20 genes enriched in Cluster 0, compared to the remainder of the cells [110, 111].

### 10x genomics analysis

The raw fastq files obtained from tumor APCs libraries were aligned to the mouse genome (mm39) using the cellranger (v. 7.0.1) pipeline. The quantification matrices were processed in the Seurat (v. 4.0.3) package for the R. Shortly, Cells containing more than 5% of their transcriptome mapped to mitochondrial reads were removed from downstream analysis. Hashtag counts and biotin antibodies were log normalized using the NormalizeData function. Cells were classified into their biological conditions based on hashtag quantification using the HTODemux function. Cells containing two or more hashtags were marked as doublets and removed. The enrichR package was used along with the Panglao database (PanglaoDB_Augmented_2021) for identifying non-APC cells. Finally, the working dataset was generated by normalizing the raw matrix of counts using the SCTransform function. We used the Wilcoxon-Rank-Test to generate differentially expressed genes with only Bonferroni adjusted-p values of 0.05 or less and log2 fold-changes of 0.3 or more were considered as significantly expressed. Signature scores were produced by running the AddModuleScore function available in Seurat. The database containing signature lists was obtained from the Molecular Signatures Database (MSigDB). Pseudo-time trajectories were calculated using the SlingShot (v. 2.7.0) package for R. Finally, to classify cells from our photo-activation experiment produced using smart-seq2 libraries into our 10x genomics experiment we used the MapQuery function available in Seurat (5.0.2).

### Statistical analysis

All statistical analyses was performed using data from at least three biological replicates, with the exact number of replicates stated in each figure legend. Unless stated otherwise, all statistical analyses were performed using Graphpad Prism v.8 software. Unpaired Student’s t test was used for most pairwise comparisons, except instances indicated in figure legends where paired Student t test or Wilcoxon signed-rank test were used. For experiments involving more than two groups of animals one-way ANOVA was used with Tukey’s test comparing the means of every treatment to the means of every other treatment. Differences with p-values < 0.05 were considered as statistically significant.

## SUPPLEMENTARY FIGURES AND LEGENDS

**Figure S1.**
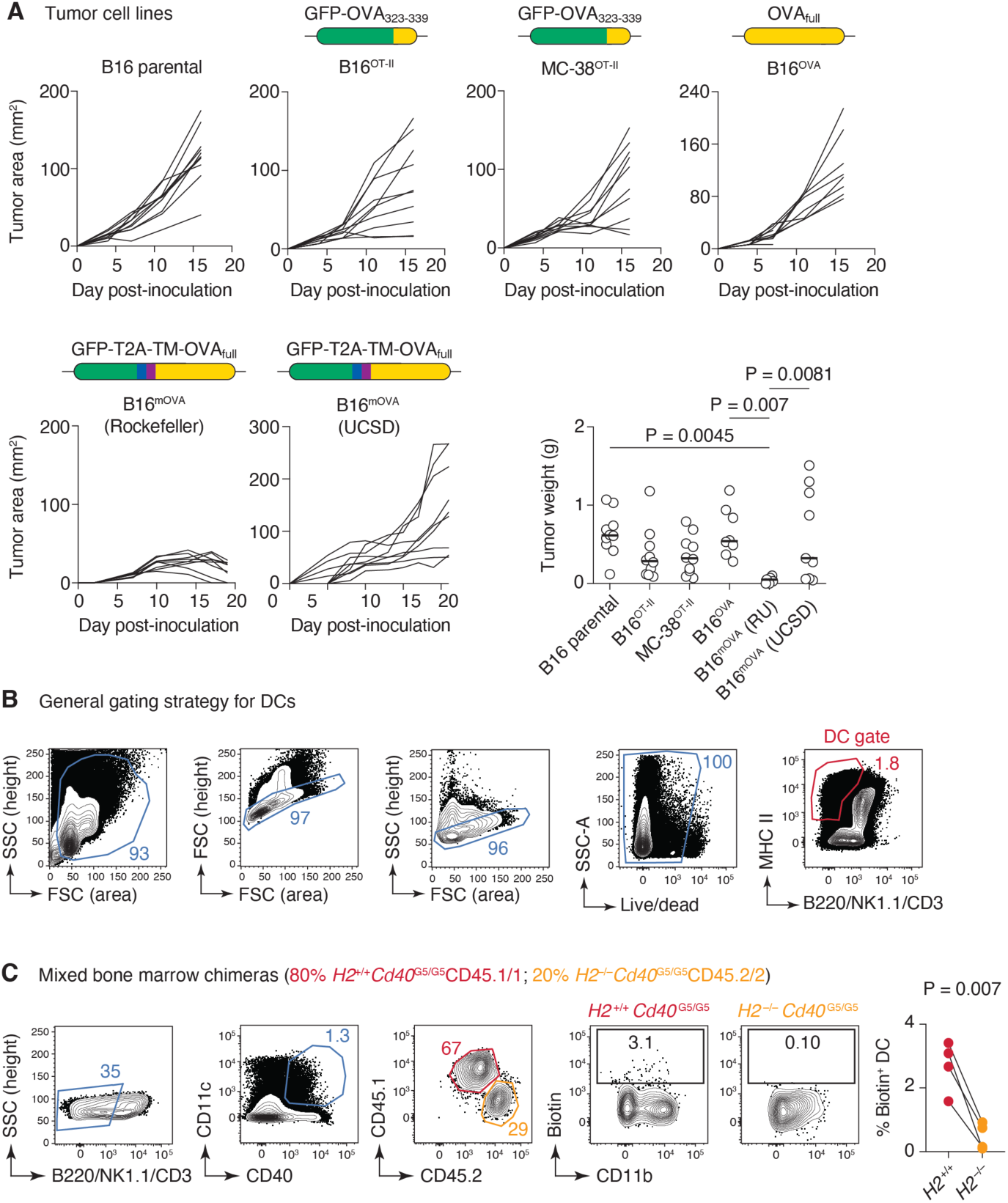
Experimental setup and additional controls. **(A)**B16 melanoma and MC-38 colon adenocarcinoma cell lines were transduced with the indicated constructs and injected into recipient mice. Tumor growth curves are shown for each indicated tumor line, compared to the previously generated parental B16 and B16^OVA^ lines. The B16mOVA line was grown in mice housed either at the Rockefeller University or at UCSD for different experiments (See Fig. 3, from which the growth and weight data are reproduced). Graph at bottom right shows tumor weight for each indicated tumor line. P-values were calculated using one-way ANOVA followed by Tukey’s multiple comparison test for the means; only p-values < 0.05 are shown. **(B)** Contour plots showing the general gating strategy for tdLN DCs prior to gating on MHC-II/CD11c. **(C)** Mixed chimeras were generated with 80% *H2*^+/+^.*Cd40*^G5/G5^ and 20% *H2*^−/−^.*Cd40*^G5/G5^ bone marrow cells transferred into lethally-irradiated WT C57BL6 mice. Contour plots show the gating strategy for MHC-II-deficient DCs and percentage of biotin^+^ DCs in tdLN in each donor population. Right, quantification of biotin^+^ DCs among MHC-II-sufficient and -deficient DCs in (n = 4 mice). P-value is for paired T-test.

**Figure S2.**
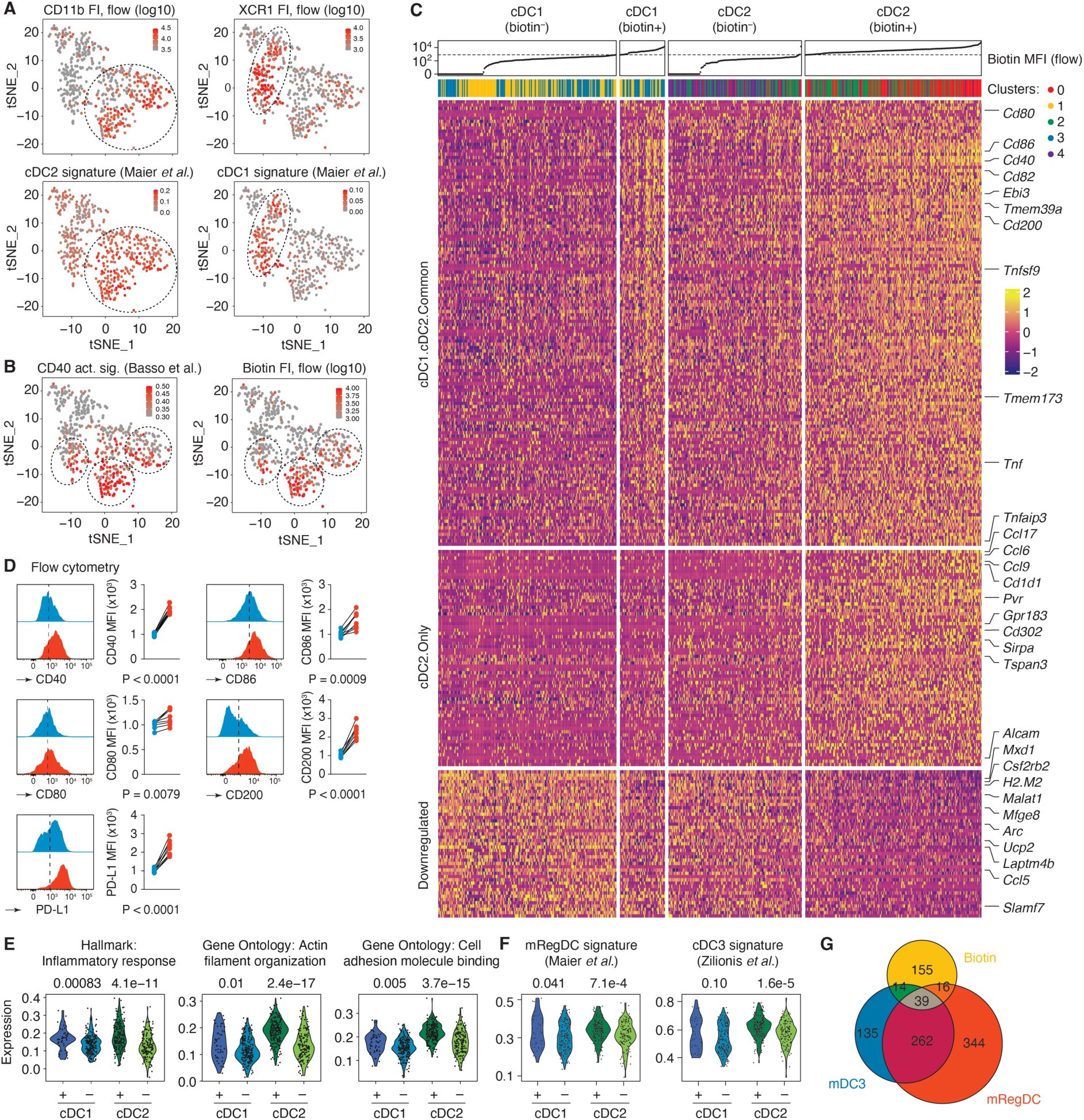
Single-cell transcriptomics of tdLN DCs. **(A)** Fluorescence intensity (FI) of CD11b and XCR1 staining (top) and expression of cDC1 and cDC2 gene expression signatures among DCs. t-SNE plot as in Fig. 2A. FI data were obtained from flow cytometry index-sorting files; cDC1 and cDC2 signatures were obtained from the literature [20]. **(B)** As in (A), showing a signature of CD40 activation in B cells [29] (left) and biotin FI (LIPSTIC signal) from index sorting (right). **(C)** Heatmap showing expression of genes significantly modulated in biotin^+^ vs. biotin^−^ DCs. The uppermost rows show biotin FI from flow cytometry, followed by transcriptional clusters as shown in Fig. 2A. Genes are sorted into 3 clusters corresponding to genes upregulated in biotin^+^ cDC1 and cDC2 (top), genes upregulated in biotin^+^ cDC2 only (middle) and genes downregulated in biotin^+^ cDC1 and cDC2 (bottom). Selected genes are indicated in the right. **(D)** Mean fluorescent intensity (MFI) normalized to the average of biotin-negative DCs for the indicated molecules. Biotin+(red) and biotin-(blue) DCs within tdLN (n = 8, from 2 independent experiments). **(E)** Violin plots showing most upregulated gene signatures in both biotin+ cDC1 and cDC2 compared to biotin– cDC1 and cDC2. **(F)** Expression of mRegDC and cDC3 signatures from the literature [20, 53] among biotin^+^ and biotin^−^ cDC1 and cDC2. **(G)** Venn diagram of the overlap between LIPSTIC^+^, cDC3, and mRegDC signatures. Paired t-test was used to measure P value for (D), Wilcoxon signed-rank test was used to measure P value for (E-G).

**Figure S3.**
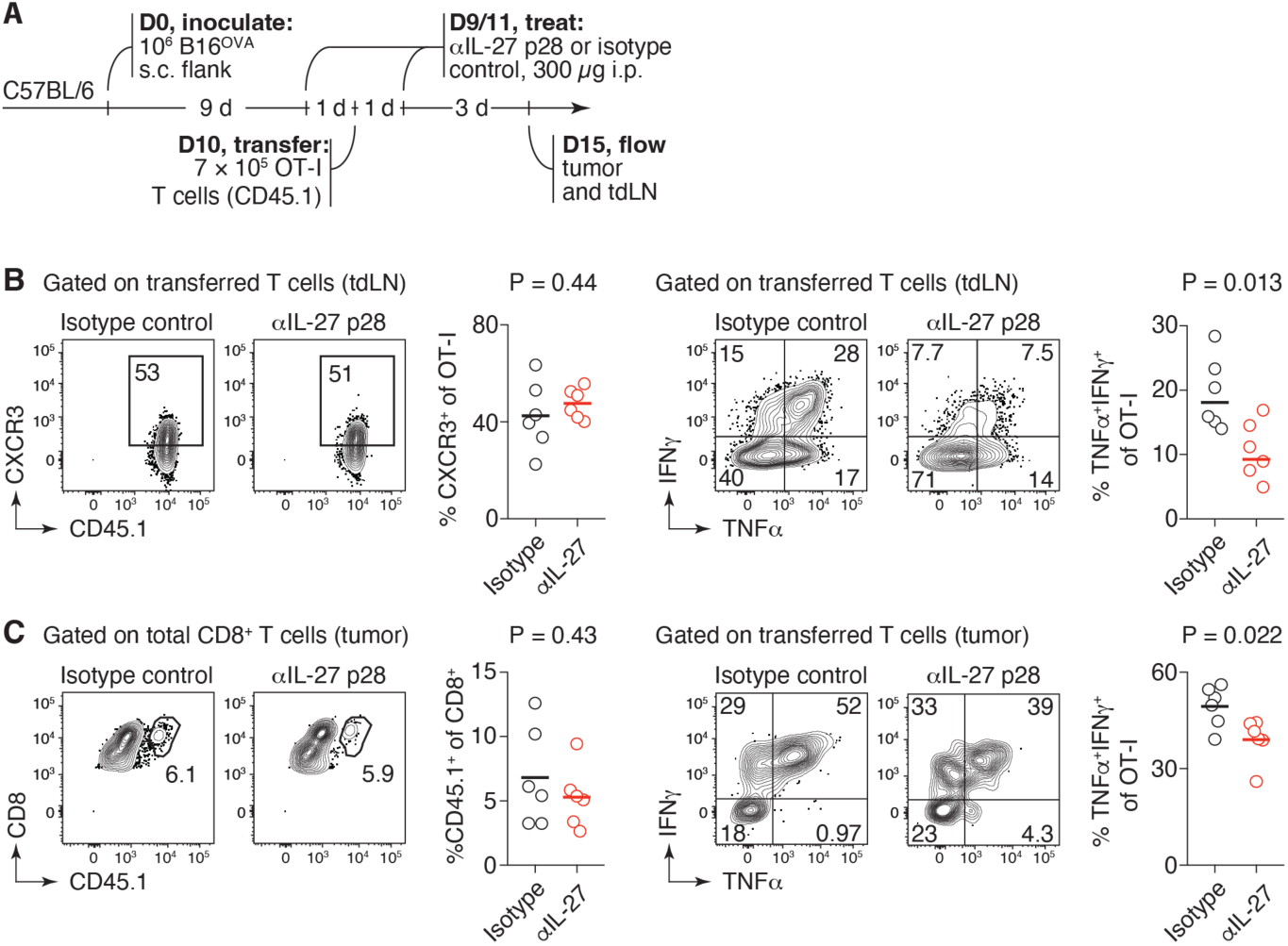
IL-27 blockade enhances CD8^+^ T cell priming. **(A)** Schematic for IL-27 (p28) blocking in mice bearing B16^OVA^ tumors. **(B)** Contour plots and quantification showing percentages of CXCR3 positive cells among transferred OT-I cells in tdLN (left). (Right) contour plots and quantification showing percentages of IFNγ^+^ TNFα^+^ among OT-I cells in tdLN. Graphs summarize data for (n = 6 mice per group). **(C)** Percentage of OT-I cells among all CD8^+^ T cells (left) and expression of TNFα and IFNγ among OT-I T cells (right) in the tumors of isotype or anti-p28-treated mice. Graphs summarize data for (n = 6 mice per group).

**Figure S4.**
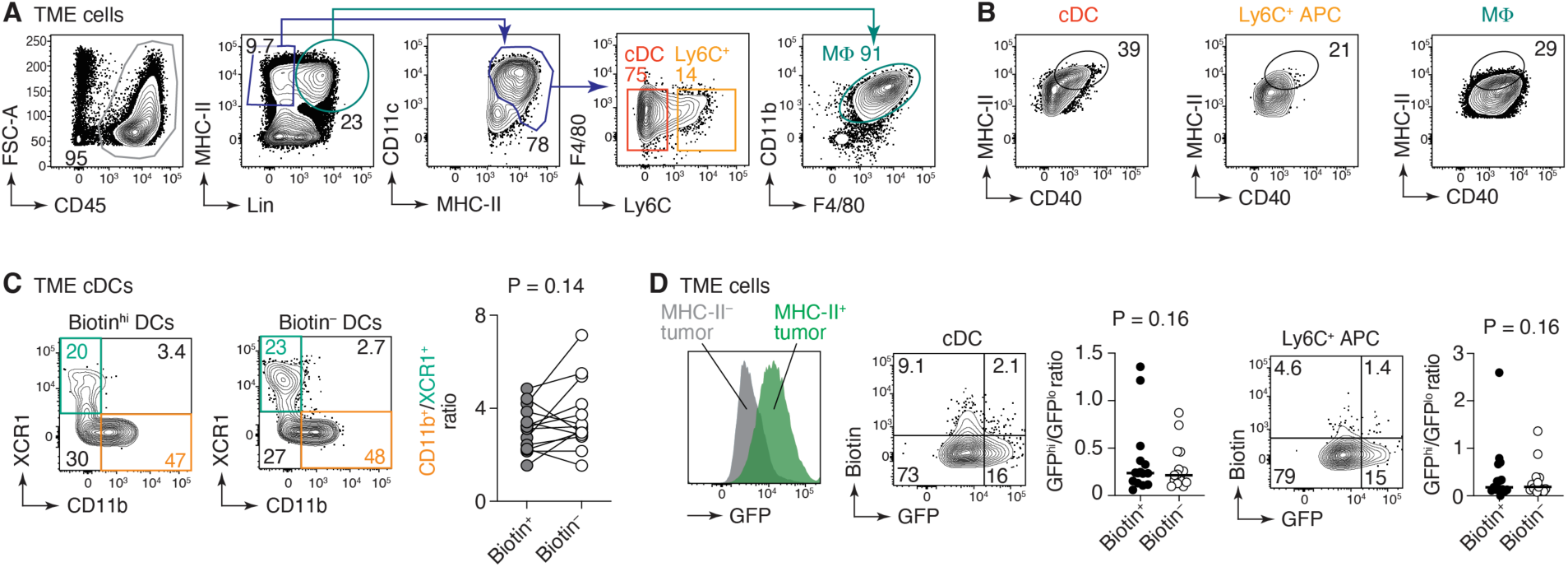
Gating strategies for APCs in tumors. **(A)** Contour plots showing how DCs, Ly6C^+^ APCs and macrophages were gated within TME. **(B)** Contour plots showing expression of CD40 and MHC-II on DCs, Ly6C^+^ APCs and macrophages. (**C**) Contour plots (left) showing percentage of cDC1 and cDC2 among biotin+ and biotin– DCs and quantification of data for (right). **(D) (**Left) Histogram plot showing GFP levels in MHC-II^-^(grey) and MHC-II^+^ cells. (Center) Contour plot and quantification showing percentage of GFP+ and biotin+DCs. (Right) Contour plot and quantification showing the percentage of GFP+ and biotin+Ly6C+APCs. Paired t-test was used for (C, D), (n = 14 mice, 4 independent experiments).

**Figure S5.**
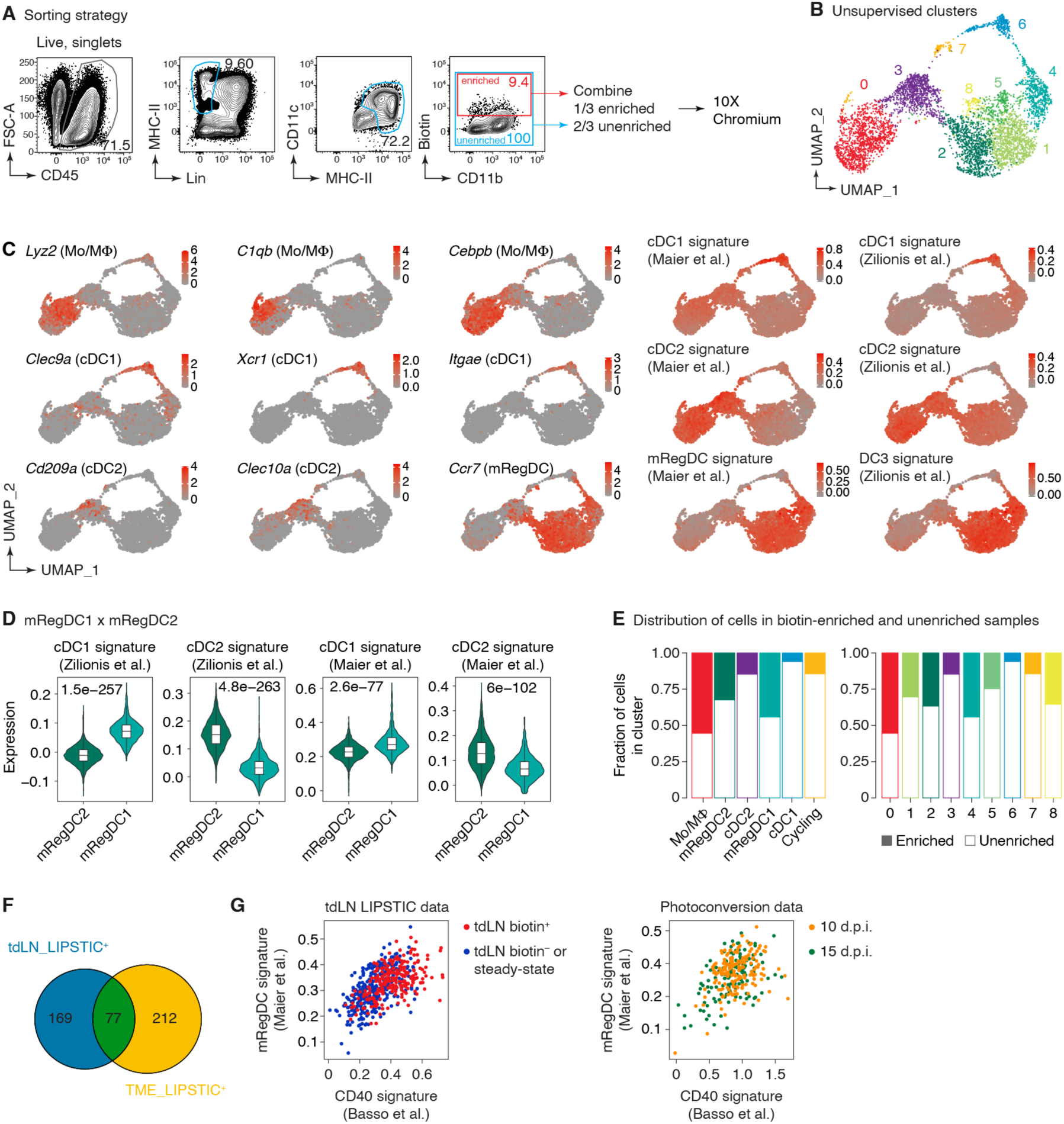
Single-cell transcriptomics of DCs in the TME. **(A)** Contour plots show sorting strategy for the 10x Genomics experiment. **(B)** UMAP plot showing unsupervised cluster distribution of APCs within TME. **(C)** UMAP plots showing the expression of landmark APC genes used for cluster annotation in Fig. 5A. Lyz2, C1qb, and *Cebpb* were used to indicate Mo/MFs; *Clec9a, Xcr1,* and *Itgae*were used to indicate cDC1s; *Cd209a, Clec10a* for cDC2s; and *Ccr7* for mRegDCs. Distribution of cDC1, cDC2, and mRegDC signatures [20, 53]. **(D)** Violin plots showing expression of cDC1 and cDC2 gene signatures [20, 53] in mRegDC1 and mRegDC2 cells. P-values are for Wilcoxon signed-rank test. **(E)** Distribution of cells in biotin-enriched and unenriched samples in annotated clusters (left) and in unsupervised clusters (right). **(F)** Venn diagram showing overlap between tdLN_LIPSTIC^+^ and TME_LIPSTIC^+^ gene signatures. (**G**) Scatter plot of CD40 and mRegDC gene signatures in tdLN DCs from Fig. 2A (left) and photoconverted DCs from Fig. 5I (right).

**Figure S6.**
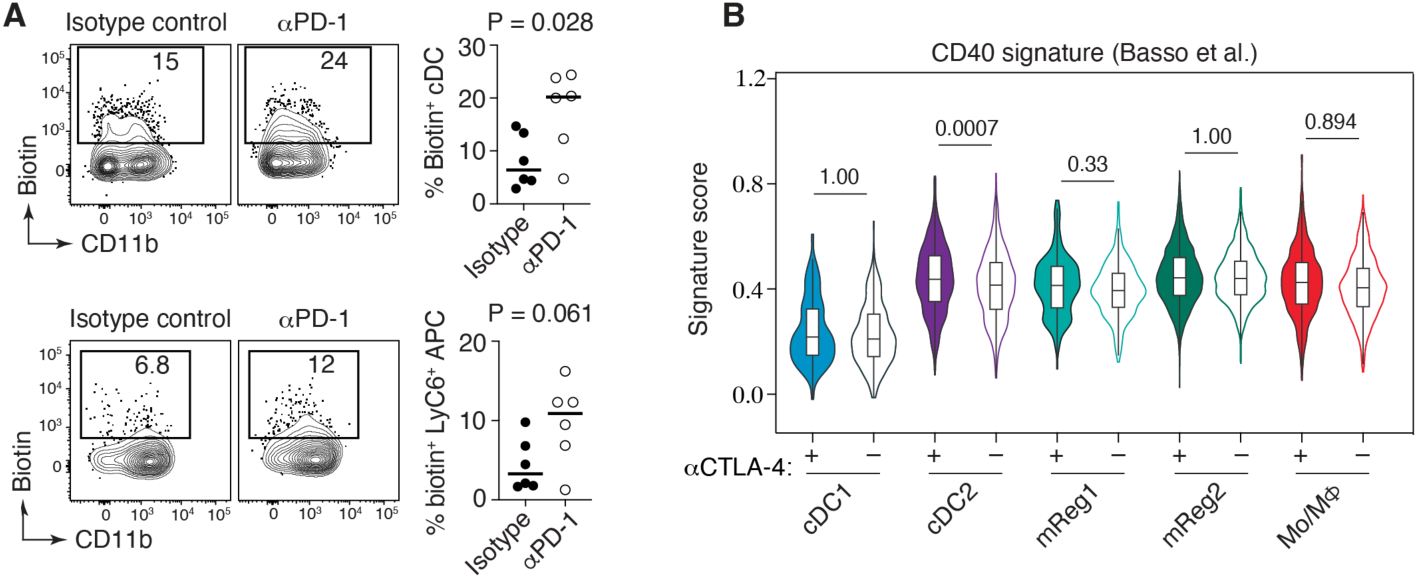
Checkpoint blockade increases APC-T cell interactions within TME. **(A)** Experimental setup as in Fig. 6.D but mice were treated with a blocking antibody to PD-1 instead of CTLA-4. Contour plots and quantification of biotin^+^ cells among DCs and Ly6C^+^ APCs in the TME as indicated (n =6 for isotype control and n = 6 for anti-PD-1, from 2 independent experiments). **(B)** Violin plots showing expression of CD40 target genes [29] in the indicated populations from anti-CTLA-4 and isotype control-treated mice.

**Table S1.** Antibodies used for flow cytometry.

| **Antibody** | **Clone** | **Cat. #** | **Vendor** |
| --- | --- | --- | --- |
| BUV 805 anti-mouse CD45 | I3/2.3 | 752415 | BD |
| BUV 496 anti-mouse CD11c | N418 | 750450 | BD |
| BUV 395 anti-mouse CD279 | J43 | 744549 | BD |
| BUV 395 anti-mouse CD45 | 30-F11 | 564279 | BD |
| BUV 395 anti-mouse CD45.1 | A20 | 565212 | BD |
| BUV 395 anti-mouse CD45.1 | A20 | 565212 | BD |
| BUV 395 anti-mouse I-A/I-E | 2G9 | 743876 | BD |
| BV 421 anti-mouse CD274 (PD-L1) | 10F.9G2 | 124315 | BioLegend |
| BV 421 anti-mouse CD279 (PD-1) | 29F.1A12 | 135221 | BioLegend |
| BV 421 anti-mouse CD4 | GK1.5 | 100438 | BioLegend |
| BV 421 anti-mouse CD64 | X54-5/7.1 | 139309 | BioLegend |
| BV 421 anti-mouse CD86 | GL-1 | 105032 | BioLegend |
| BV 421 anti-mouse I-A/I-E | M5/114.15.2 | 107632 | BioLegend |
| BV 421 anti-mouse CD40 | 3/23' | 562846 | BD |
| BV 605 anti-mouse F4/80 | T45-2342 | 743281 | BD |
| BV 605 anti-mouse CD45.2 | 104 | 109841 | BioLegend |
| BV 605 anti-mouse CD86 | GL-1 | 105037 | BioLegend |
| BV 605 anti-mouse CD8 | 53-6.7 | 100744 | BioLegend |
| BV 711 anti-mouse CD3ε | 145-2 C11 | 100349 | BioLegend |
| BV 711 anti-mouse CD4 | RM4-5 | 100557 | BioLegend |
| BV 711 anti-mouse/human CD11b | M1/70 | 101242 | BioLegend |
| BV 785 anti-mouse CD3 | 17A2 | 100232 | BioLegend |
| BV 785 anti-mouse NK-1.1 | PK136 | 108749 | BioLegend |
| BV 785 anti-mouse/human CD45R/B220 | RA3-6B2 | 103246 | BioLegend |
| PE anti-biotin | 1D4-C5 | 409004 | BioLegend |
| PE anti-biotin | Bio3-18E7 | 130-113-291 | Miltenyi |
| PE anti-mouse CD183 (CXCR3) | CXCR3-173 | 126505 | BioLegend |
| PE anti-mouse TNF-α | MP6-XT22 | 506306 | BioLegend |
| PE/Cy7 anti-mouse CD223 (LAG-3) | C9B7W | 125226 | BioLegend |
| PE/Cy7 anti-mouse CD45.1 | A20 | 110730 | BioLegend |
| PerCP-eFluor 710 TIGIT | GIGD7 | 46-9501-80 | Invitrogen |
| PerCP-eFluor™ 710 anti-mouse CD200 | OX90 | 46-5200-82 | ThermoFisher |
| PerCP/Cy5.5 anti-mouse CD80 | 16-10A1 | 104722 | BioLegend |
| PerCP/Cy5.5 anti-mouse/rat XCR1 | ZET | 148208 | BioLegend |
| APC anti-mouse Ly-6C | HK1.4 | 128016 | BioLegend |
| APC anti-mouse CD11c | N418 | 117310 | BioLegend |
| APC anti-mouse CD64 | X54-5/7.1 | 17-0641-82 | ThermoFisher |
| APC anti-mouse IFN | XMG1.2 | 17-7311-82 | ThermoFisher |
| TotalSeq-C0301 Hashtag 1 | 30-F11& M1/42 | 155861 | BioLegend |
| TotalSeq-C0302 Hashtag 2 | 30-F11&M1/42 | 155863 | BioLegend |
| TotalSeq-C0303 Hashtag 3 | 30-F11& M1/42 | 155865 | BioLegend |
| TotalSeq-C0304 Hashtag 4 | 30-F11& M1/42 | 155867 | BioLegend |
| TotalSeq-C0305 Hashtag 5 | 30-F11& M1/42 | 155869 | BioLegend |
| TotalSeq-C0306 Hashtag 6 | 30-F11& M1/42 | 155871 | BioLegend |
| TotalSeq-C0307 Hashtag 7 | 30-F11& M1/42 | 155873 | BioLegend |
| TotalSeq-C0308 Hashtag 8 | 30-F11& M1/42 | 155875 | BioLegend |
| TotalSeq-C0309 Hashtag 9 | 30-F11& M1/42 | 155877 | BioLegend |
| TotalSeq-C0310 Hashtag 10 | 30-F11& M1/42 | 155879 | BioLegend |
| TotalSeq-C0306 Hashtag 11 | 30-F11& M1/42 | 155881 | BioLegend |
| TotalSeq-C0306 Hashtag 12 | 30-F11& M1/42 | 155883 | BioLegend |
| TotalSeq-C0306 Hashtag 13 | 30-F11& M1/42 | 155885 | BioLegend |
| TotalSeq-C0306 Hashtag 14 | 30-F11& M1/42 | 155887 | BioLegend |
| TotalSeq-C0436 anti-biotin | 1D4-C5 | 409011 | BioLegend |

## References

1. Leach, D.R., M.F. Krummel, and J.P. Allison, Enhancement of antitumor immunity by CTLA-4 blockade. Science, 1996. 271(5256): p. 1734–6.

2. Sharma, P. and J.P. Allison, Immune checkpoint targeting in cancer therapy: toward combination strategies with curative potential. Cell, 2015. 161(2): p. 205–14.

3. DeVita, V.T., Jr. and S.A. Rosenberg, Two hundred years of cancer research. N Engl J Med, 2012. 366(23): p. 2207–14.

4. Chudnovskiy, A., G. Pasqual, and G.D. Victora, Studying interactions between dendritic cells and T cells in vivo. Curr Opin Immunol, 2019. 58: p. 24–30.

5. Merad, M., et al., The dendritic cell lineage: ontogeny and function of dendritic cells and their subsets in the steady state and the inflamed setting. Annu Rev Immunol, 2013. 31: p. 563–604.

6. Wculek, S.K., et al., Dendritic cells in cancer immunology and immunotherapy. Nat Rev Immunol, 2020. 20(1): p. 7–24.

7. Ruhland, M.K., et al., Visualizing Synaptic Transfer of Tumor Antigens among Dendritic Cells. Cancer Cell, 2020. 37(6): p. 786–799 e5.

8. Murphy, T.L., et al., Transcriptional Control of Dendritic Cell Development. Annu Rev Immunol, 2016. 34: p. 93–119.

9. Hildner, K., et al., Batf3 deficiency reveals a critical role for CD8alpha+ dendritic cells in cytotoxic T cell immunity. Science, 2008. 322(5904): p. 1097–100.

10. Roberts, E.W., et al., Critical Role for CD103(+)/CD141(+) Dendritic Cells Bearing CCR7 for Tumor Antigen Trafficking and Priming of T Cell Immunity in Melanoma. Cancer Cell, 2016. 30(2): p. 324–336.

11. Salmon, H., et al., Expansion and Activation of CD103(+) Dendritic Cell Progenitors at the Tumor Site Enhances Tumor Responses to Therapeutic PD-L1 and BRAF Inhibition. Immunity, 2016. 44(4): p. 924–38.

12. Ferris, S.T., et al., cDC1 prime and are licensed by CD4(+) T cells to induce anti-tumour immunity. Nature, 2020. 584(7822): p. 624–629.

13. Binnewies, M., et al., Unleashing Type-2 Dendritic Cells to Drive Protective Antitumor CD4(+) T Cell Immunity. Cell, 2019. 177(3): p. 556–571 e16.

14. Broz, M.L., et al., Dissecting the Tumor Myeloid Compartment Reveals Rare Activating Antigen-Presenting Cells Critical for T Cell Immunity. Cancer Cell, 2014. 26(6): p. 938.

15. Sanchez-Paulete, A.R., et al., Cancer Immunotherapy with Immunomodulatory Anti-CD137 and Anti-PD-1 Monoclonal Antibodies Requires BATF3-Dependent Dendritic Cells. Cancer Discov, 2016. 6(1): p. 71–9.

16. Spranger, S., R. Bao, and T.F. Gajewski, Melanoma-intrinsic beta-catenin signalling prevents anti-tumour immunity. Nature, 2015. 523(7559): p. 231–5.

17. Spranger, S., et al., Tumor-Residing Batf3 Dendritic Cells Are Required for Effector T Cell Trafficking and Adoptive T Cell Therapy. Cancer Cell, 2017. 31(5): p. 711–723 e4.

18. Marangoni, F., et al., Expansion of tumor-associated Treg cells upon disruption of a CTLA-4-dependent feedback loop. Cell, 2021. 184(15): p. 3998–4015 e19.

19. Garris, C.S., et al., Successful Anti-PD-1 Cancer Immunotherapy Requires T Cell-Dendritic Cell Crosstalk Involving the Cytokines IFN-gamma and IL-12. Immunity, 2018. 49(6): p. 1148–1161 e7.

20. Maier, B., et al., A conserved dendritic-cell regulatory program limits antitumour immunity. Nature, 2020. 580(7802): p. 257–262.

21. Villani, A.C., et al., Single-cell RNA-seq reveals new types of human blood dendritic cells, monocytes, and progenitors. Science, 2017. 356(6335).

22. Dutertre, C.A., et al., Single-Cell Analysis of Human Mononuclear Phagocytes Reveals Subset-Defining Markers and Identifies Circulating Inflammatory Dendritic Cells. Immunity, 2019. 51(3): p. 573–589 e8.

23. Ginhoux, F., M. Guilliams, and M. Merad, Expanding dendritic cell nomenclature in the single-cell era. Nat Rev Immunol, 2022. 22(2): p. 67–68.

24. Heras-Murillo, I., et al., Dendritic cells as orchestrators of anticancer immunity and immunotherapy. Nat Rev Clin Oncol, 2024. 21(4): p. 257–277.

25. Pittet, M.J., et al., Dendritic cells as shepherds of T cell immunity in cancer. Immunity, 2023. 56(10): p. 2218–2230.

26. Del Prete, A., et al., Dendritic cell subsets in cancer immunity and tumor antigen sensing. Cell Mol Immunol, 2023. 20(5): p. 432–447.

27. Chen, D.S. and I. Mellman, Oncology meets immunology: the cancer-immunity cycle. Immunity, 2013. 39(1): p. 1–10.

28. Pasqual, G., et al., Monitoring T cell-dendritic cell interactions in vivo by intercellular enzymatic labelling. Nature, 2018. 553(7689): p. 496–500.

29. Basso, K., et al., Tracking CD40 signaling during germinal center development. Blood, 2004. 104(13): p. 4088–96.

30. Ghosh, S. and M.S. Hayden, New regulators of NF-kappaB in inflammation. Nat Rev Immunol, 2008. 8(11): p. 837–48.

31. Lee, E.G., et al., Failure to regulate TNF-induced NF-kappaB and cell death responses in A20-deficient mice. Science, 2000. 289(5488): p. 2350–4.

32. Peng, Q., et al., PD-L1 on dendritic cells attenuates T cell activation and regulates response to immune checkpoint blockade. Nat Commun, 2020. 11(1): p. 4835.

33. Rossjohn, J., et al., Recognition of CD1d-restricted antigens by natural killer T cells. Nat Rev Immunol, 2012. 12(12): p. 845–57.

34. Zimmerman, A.W., et al., Long-term engagement of CD6 and ALCAM is essential for T-cell proliferation induced by dendritic cells. Blood, 2006. 107(8): p. 3212–20.

35. Kumanogoh, A., et al., Nonredundant roles of Sema4A in the immune system: defective T cell priming and Th1/Th2 regulation in Sema4A-deficient mice. Immunity, 2005. 22(3): p. 305–16.

36. Groettrup, M., C.J. Kirk, and M. Basler, Proteasomes in immune cells: more than peptide producers? Nat Rev Immunol, 2010. 10(1): p. 73–8.

37. Everitt, A.R., et al., IFITM3 restricts the morbidity and mortality associated with influenza. Nature, 2012. 484(7395): p. 519–23.

38. Tran, Q., et al., TMEM39A and Human Diseases: A Brief Review. Toxicol Res, 2017. 33(3): p. 205–209.

39. Decout, A., et al., The cGAS-STING pathway as a therapeutic target in inflammatory diseases. Nat Rev Immunol, 2021. 21(9): p. 548–569.

40. Zhao, X., et al., CCL9 is secreted by the follicle-associated epithelium and recruits dome region Peyer’s patch CD11b+ dendritic cells. J Immunol, 2003. 171(6): p. 2797–803.

41. Semmling, V., et al., Alternative cross-priming through CCL17-CCR4-mediated attraction of CTLs toward NKT cell-licensed DCs. Nat Immunol, 2010. 11(4): p. 313–20.

42. Coelho, A.L., et al., The chemokine CCL6 promotes innate immunity via immune cell activation and recruitment. J Immunol, 2007. 179(8): p. 5474–82.

43. Geijtenbeek, T.B. and S.I. Gringhuis, C-type lectin receptors in the control of T helper cell differentiation. Nat Rev Immunol, 2016. 16(7): p. 433–48.

44. Wynn, T.A., Type 2 cytokines: mechanisms and therapeutic strategies. Nat Rev Immunol, 2015. 15(5): p. 271–82.

45. Gatto, D., et al., The chemotactic receptor EBI2 regulates the homeostasis, localization and immunological function of splenic dendritic cells. Nat Immunol, 2013. 14(5): p. 446–53.

46. Li, J., et al., EBI2 augments Tfh cell fate by promoting interaction with IL-2-quenching dendritic cells. Nature, 2016. 533(7601): p. 110–4.

47. Yi, T. and J.G. Cyster, EBI2-mediated bridging channel positioning supports splenic dendritic cell homeostasis and particulate antigen capture. Elife, 2013. 2: p. e00757.

48. Brown, C.C., et al., Transcriptional Basis of Mouse and Human Dendritic Cell Heterogeneity. Cell, 2019. 179(4): p. 846–863 e24.

49. Jinushi, M., et al., MFG-E8-mediated uptake of apoptotic cells by APCs links the pro- and antiinflammatory activities of GM-CSF. J Clin Invest, 2007. 117(7): p. 1902–13.

50. Jeon, M.S., et al., Essential role of the E3 ubiquitin ligase Cbl-b in T cell anergy induction. Immunity, 2004. 21(2): p. 167–77.

51. Liberzon, A., et al., The Molecular Signatures Database (MSigDB) hallmark gene set collection. Cell Syst, 2015. 1(6): p. 417–425.

52. Subramanian, A., et al., Gene set enrichment analysis: a knowledge-based approach for interpreting genome-wide expression profiles. Proc Natl Acad Sci U S A, 2005. 102(43): p. 15545–50.

53. Zilionis, R., et al., Single-Cell Transcriptomics of Human and Mouse Lung Cancers Reveals Conserved Myeloid Populations across Individuals and Species. Immunity, 2019. 50(5): p. 1317–1334 e10.

54. Corbett, T.H., et al., Tumor induction relationships in development of transplantable cancers of the colon in mice for chemotherapy assays, with a note on carcinogen structure. Cancer Res, 1975. 35(9): p. 2434–9.

55. Yoshida, H. and C.A. Hunter, The immunobiology of interleukin-27. Annu Rev Immunol, 2015. 33: p. 417–43.

56. Fabbi, M., G. Carbotti, and S. Ferrini, Dual Roles of IL-27 in Cancer Biology and Immunotherapy. Mediators Inflamm, 2017. 2017: p. 3958069.

57. Chihara, N., et al., Induction and transcriptional regulation of the co-inhibitory gene module in T cells. Nature, 2018. 558(7710): p. 454–459.

58. Zhu, J., et al., IL-27 gene therapy induces depletion of Tregs and enhances the efficacy of cancer immunotherapy. JCI Insight, 2018. 3(7).

59. Groom, J.R., et al., CXCR3 chemokine receptor-ligand interactions in the lymph node optimize CD4+ T helper 1 cell differentiation. Immunity, 2012. 37(6): p. 1091–103.

60. Mikucki, M.E., et al., Non-redundant requirement for CXCR3 signalling during tumoricidal T-cell trafficking across tumour vascular checkpoints. Nat Commun, 2015. 6: p. 7458.

61. Magen, A., et al., Intratumoral dendritic cell-CD4(+) T helper cell niches enable CD8(+) T cell differentiation following PD-1 blockade in hepatocellular carcinoma. Nat Med, 2023. 29(6): p. 1389–1399.

62. Nakandakari-Higa, S., et al., Universal recording of immune cell interactions in vivo. Nature, 2024. 627(8003): p. 399–406.

63. Rizzitelli, A., et al., Serpinb9 (Spi6)-deficient mice are impaired in dendritic cell-mediated antigen cross-presentation. Immunol Cell Biol, 2012. 90(9): p. 841–51.

64. Pishesha, N., T.J. Harmand, and H.L. Ploegh, A guide to antigen processing and presentation. Nat Rev Immunol, 2022. 22(12): p. 751–764.

65. Arron, J.R., et al., Regulation of the subcellular localization of tumor necrosis factor receptor-associated factor (TRAF)2 by TRAF1 reveals mechanisms of TRAF2 signaling. J Exp Med, 2002. 196(7): p. 923–34.

66. Donninelli, G., et al., Dual requirement for STAT signaling in dendritic cell immunobiology. Immunobiology, 2018. 223(3): p. 342–347.

67. Chen, L. and D.B. Flies, Molecular mechanisms of T cell co-stimulation and co-inhibition. Nat Rev Immunol, 2013. 13(4): p. 227–42.

68. Di Pilato, M., et al., CXCR6 positions cytotoxic T cells to receive critical survival signals in the tumor microenvironment. Cell, 2021. 184(17): p. 4512–4530 e22.

69. Wei, J., et al., Critical role of dendritic cell-derived IL-27 in antitumor immunity through regulating the recruitment and activation of NK and NKT cells. J Immunol, 2013. 191(1): p. 500–8.

70. Kitano, M., et al., Imaging of the cross-presenting dendritic cell subsets in the skin-draining lymph node. Proc Natl Acad Sci U S A, 2016. 113(4): p. 1044–9.

71. Nowotschin, S. and A.K. Hadjantonakis, Use of KikGR a photoconvertible green-to-red fluorescent protein for cell labeling and lineage analysis in ES cells and mouse embryos. BMC Dev Biol, 2009. 9: p. 49.

72. Tsutsui, H., et al., Semi-rational engineering of a coral fluorescent protein into an efficient highlighter. EMBO Rep, 2005. 6(3): p. 233–8.

73. Lee, C.Y.C., et al., Tumour-retained activated CCR7(+) dendritic cells are heterogeneous and regulate local anti-tumour cytolytic activity. Nat Commun, 2024. 15(1): p. 682.

74. Mellman, I., et al., The cancer-immunity cycle: Indication, genotype, and immunotype. Immunity, 2023. 56(10): p. 2188–2205.

75. Pardoll, D.M., The blockade of immune checkpoints in cancer immunotherapy. Nat Rev Cancer, 2012. 12(4): p. 252–64.

76. Chen, B., et al., Unraveling the Heterogeneity and Ontogeny of Dendritic Cells Using Single-Cell RNA Sequencing. Front Immunol, 2021. 12: p. 711329.

77. Esterhazy, D., et al., Classical dendritic cells are required for dietary antigen-mediated induction of peripheral T(reg) cells and tolerance. Nat Immunol, 2016. 17(5): p. 545–55.

78. Giladi, A., et al., Dissecting cellular crosstalk by sequencing physically interacting cells. Nat Biotechnol, 2020. 38(5): p. 629–637.

79. Altman, J.D., et al., Phenotypic analysis of antigen-specific T lymphocytes. Science, 1996. 274(5284): p. 94–6.

80. Krishnamurty, A.T., et al., Somatically Hypermutated Plasmodium-Specific IgM(+) Memory B Cells Are Rapid, Plastic, Early Responders upon Malaria Rechallenge. Immunity, 2016. 45(2): p. 402–14.

81. Cohen, M., et al., The interaction of CD4(+) helper T cells with dendritic cells shapes the tumor microenvironment and immune checkpoint blockade response. Nat Cancer, 2022. 3(3): p. 303–317.

82. Lesley, R., et al., Naive CD4 T cells constitutively express CD40L and augment autoreactive B cell survival. Proc Natl Acad Sci U S A, 2006. 103(28): p. 10717–22.

83. Grewal, I.S. and R.A. Flavell, The role of CD40 ligand in costimulation and T-cell activation. Immunol Rev, 1996. 153: p. 85–106.

84. Dohler, A., et al., RelB(+) Steady-State Migratory Dendritic Cells Control the Peripheral Pool of the Natural Foxp3(+) Regulatory T Cells. Front Immunol, 2017. 8: p. 726.

85. Borst, J., et al., CD4(+) T cell help in cancer immunology and immunotherapy. Nat Rev Immunol, 2018. 18(10): p. 635–647.

86. Bennett, S.R., et al., Help for cytotoxic-T-cell responses is mediated by CD40 signalling. Nature, 1998. 393(6684): p. 478–80.

87. Schoenberger, S.P., et al., T-cell help for cytotoxic T lymphocytes is mediated by CD40-CD40L interactions. Nature, 1998. 393(6684): p. 480–3.

88. Ridge, J.P., F. Di Rosa, and P. Matzinger, A conditioned dendritic cell can be a temporal bridge between a CD4+ T-helper and a T-killer cell. Nature, 1998. 393(6684): p. 474–8.

89. Banchereau, J. and R.M. Steinman, Dendritic cells and the control of immunity. Nature, 1998. 392(6673): p. 245–52.

90. Mellman, I. and R.M. Steinman, Dendritic cells: specialized and regulated antigen processing machines. Cell, 2001. 106(3): p. 255–8.

91. Oliveira, M.M.S. and L.S. Westerberg, Cytoskeletal regulation of dendritic cells: An intricate balance between migration and presentation for tumor therapy. J Leukoc Biol, 2020. 108(4): p. 1051–1065.

92. Castellino, F., et al., Chemokines enhance immunity by guiding naive CD8+ T cells to sites of CD4+ T cell-dendritic cell interaction. Nature, 2006. 440(7086): p. 890–5.

93. Murugaiyan, G. and B. Saha, IL-27 in tumor immunity and immunotherapy. Trends Mol Med, 2013. 19(2): p. 108–16.

94. Hu, A., et al., Intra-Tumoral Delivery of IL-27 Using Adeno-Associated Virus Stimulates Anti-tumor Immunity and Enhances the Efficacy of Immunotherapy. Front Cell Dev Biol, 2020. 8: p. 210.

95. Kilgore, A.M., et al., IL-27p28 Production by XCR1(+) Dendritic Cells and Monocytes Effectively Predicts Adjuvant-Elicited CD8(+) T Cell Responses. Immunohorizons, 2018. 2(1): p. 1–11.

96. Pennock, N.D., L. Gapin, and R.M. Kedl, IL-27 is required for shaping the magnitude, affinity distribution, and memory of T cells responding to subunit immunization. Proc Natl Acad Sci U S A, 2014. 111(46): p. 16472–7.

97. Gerhard, G.M., et al., Tumor-infiltrating dendritic cell states are conserved across solid human cancers. J Exp Med, 2021. 218(1).

98. Sharma, P. and J.P. Allison, Dissecting the mechanisms of immune checkpoint therapy. Nat Rev Immunol, 2020. 20(2): p. 75–76.

99. Madsen, L., et al., Mice lacking all conventional MHC class II genes. Proc Natl Acad Sci U S A, 1999. 96(18): p. 10338–43.

100. Lee, P.P., et al., A critical role for Dnmt1 and DNA methylation in T cell development, function, and survival. Immunity, 2001. 15(5): p. 763–74.

101. Hogquist, K.A., et al., T cell receptor antagonist peptides induce positive selection. Cell, 1994. 76(1): p. 17–27.

102. Barnden, M.J., et al., Defective TCR expression in transgenic mice constructed using cDNA-based alpha- and beta-chain genes under the control of heterologous regulatory elements. Immunol Cell Biol, 1998. 76(1): p. 34–40.

103. Caton, M.L., M.R. Smith-Raska, and B. Reizis, Notch-RBP-J signaling controls the homeostasis of CD8-dendritic cells in the spleen. J Exp Med, 2007. 204(7): p. 1653–64.

104. Quadros, R.M., et al., Easi-CRISPR: a robust method for one-step generation of mice carrying conditional and insertion alleles using long ssDNA donors and CRISPR ribonucleoproteins. Genome Biol, 2017. 18(1): p. 92.

105. Engels, B., et al., Retroviral vectors for high-level transgene expression in T lymphocytes. Hum Gene Ther, 2003. 14(12): p. 1155–68.

106. Szymczak, A.L. and D.A. Vignali, Development of 2A peptide-based strategies in the design of multicistronic vectors. Expert Opin Biol Ther, 2005. 5(5): p. 627–38.

107. Trombetta, J.J., et al., Preparation of Single-Cell RNA-Seq Libraries for Next Generation Sequencing. Curr Protoc Mol Biol, 2014. 107: p. 4 22 1-4 22 17.

108. Hao, Y., et al., Integrated analysis of multimodal single-cell data. Cell, 2021. 184(13): p. 3573–3587 e29.

109. Korotkevich, G., et al., 2021.

110. Stephens, M., False discovery rates: a new deal. Biostatistics, 2017. 18(2): p. 275–294.

111. Love, M.I., W. Huber, and S. Anders, Moderated estimation of fold change and dispersion for RNA-seq data with DESeq2. Genome Biol, 2014. 15(12): p. 550.

